# DMRU: Generative Deep-Learning to unravel condition specific cytosine methylation in plants

**DOI:** 10.1101/2025.02.06.635186

**Authors:** Sagar Gupta, Anchit Kumar, Veerbhan Kesarwani, Umesh Bhati, Ravi Shankar

## Abstract

Methylation at cytosines in plants influence spatio-temporal gene expression by regulating chromatin structure and accessibility. Some algorithms have been developed to identify DNA methylation but none of them are capable to tell the condition specific DNA methylation, making them hardly of any use. Here, we report a first of its kind an explainable Deep Encoders-Decoders generative system, DMRU, which learns the relationship between transcritpome status and DNA methylation states at any given time. It was also found that GC similarity is more relevant to the specificity of DNA methylation patterns than homology, concurring with reports of direct involvement of GC content in providing regulatory switches for DNA accessibility. Leveraging on which DMRU could perform with same level of accuracy in cross-species universal manner. In a comprehensive testing and benchmarking study across a huge volume of experimental data covering 85 different conditions, and multiple plant species, it has consistently achieved >90% accuracy. With this all, DMRU brings a completely new chapter in methylated cytosine discovery, giving a strong alternative to costly bisulfite sequencing experiments. DMRU may prove critical turning point in plant regulatory research and its acceleration.

## Introduction

DNA methylation is a key epigenetic mechanism through which spatio-temporal regulation is achieved in plants **[1]**. It involves the addition of methyl groups to specific DNA nucleotides (adenine and cytosine), primarily targeting cytosine residues in plants. The most common form of this modification is the conversion of cytosine to 5-methylcytosine (5mC) **[2]**. This modification is not static; it dynamically responds to a multitude of factors, including various environmental and developmental influences. These factors can lead to changes in the methylation landscape, affecting how genes are expressed.

DNA methylation can be investigated through advanced techniques like next-generation sequencing (NGS). Among these, whole-genome bisulfite sequencing (WGBS) **[3]** and reduced-representation bisulfite sequencing (RRBS) **[4]** stand out as the most powerful methods. WGBS provides a genome-wide view of DNA methylation, using sodium bisulfite which converts unmethylated cytosines to uracils while leaving methylated cytosines unaffected **(Figure 1)**. RRBS is a targeted, cost-effective approach focusing on CpG-rich regions, offering high-resolution data for specific loci to study regulatory elements and traits.

**Figure 1:**
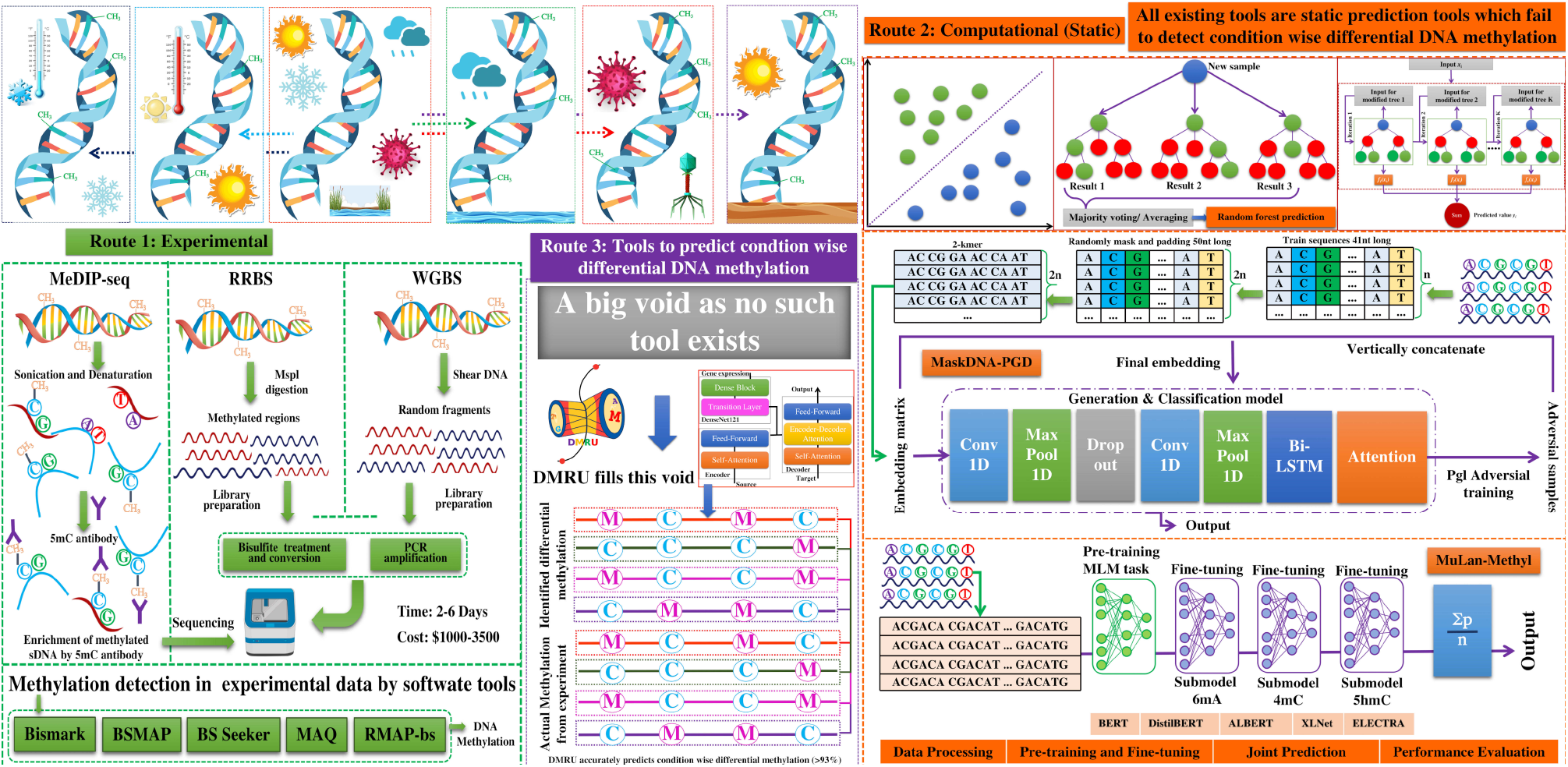
The general introduction of the problem and solution. Regulatory DNA methylations in plants regulate downstream gene’s expression in condition specific manner as DNA methylation is also condition specific and not static. The DNA methylation can be decoded through three ways: **a)** Bisulfite like experiments and subsequent sequencing and subsequent detection by software, **b)** Software tools directly considering the DNA sequence only and making static prediction of DNA methylation which completely fails to decode the condition specific DNA methylation, and **c)** Software to detect condition specific DNA-methylation. To this date, no other software solution existed to do this. DMRU, the approach presented here is the first of this kind ever. It uses information from RNA-seq transcriptome as well as target DNA sequence environment to build dynamic model of DNA methylation capable to detect condition specific differential methylation pattern. It has incorporated a DenseNet-Transformers based composite deep-learning encoder decoder system to generate the methylated states while detecting the differential condition information from the RNA-seq data.

While significant progress has been made in understanding DNA methylation in animal models, particularly in humans, there is a striking disparity in plant research. The plant kingdom comprises a vast number of species, each with unique genetic and epigenetic landscapes. Defining their wide range of habitats and survival responses. However, only a limited number of plant species have been sequenced for their DNA methylation states. In fact, fewer than 10 species have been studied, highlighting a considerable gap in our knowledge about plant methylation dynamics which is also mirrored by the minuscule development of computational tools and software resources **(Supplementary Table S1)**. This represents a stark contrast to the extensive research conducted in human. This lack of resources hampers the ability to properly explore the DNA methylations and their impact in various conditions in the plant kingdom. As research continues to evolve, there is a pressing need to bridge this gap to foster proper understanding of DNA methylation in plants and its implications for biology and agriculture.

The detection of DNA methylation has traditionally relied on experimental techniques which are both expensive and time-consuming **[5]**. WGBS costs varies significantly depending upon the service provider, sample volume, specific project requirements, and can range between approx $2,500 to $3,300 per sample. Additionally, methylation patterns varies significantly across different species, which further complicates the matter. Above all, very much like the transcriptome profiling, DNA methylome profile has unlimited possible states depending upon the condition which makes it quite unpragmatic to reveal all of them experimentally. Therefore, it has become an essentiality to develop some efficient computational approach to address these issues, a tool which could accurately profile DNA methylation profile for any given condition. To address these issues, researchers are increasingly turning towards computational approaches, particularly those based on machine learning (ML). These methods claim good accuracy. They usually classify DNA methylation by framing the identification of methylation sites as a binary classification problem. In this context, ML models are trained to differentiate between true methylation sites and non-methylation sites based on genomic sequences.

While the goal remains the same for these tools, the approaches taken differ significantly in terms of the features selected and the architectures employed. ML methods focus on manually engineered features, requiring expert knowledge and prior experience. These features, including sequence encoding and biochemical characteristics like nucleotide composition or methylation patterns **[6-7]**, can limit the adaptability of the models across species or contexts due to varying biological backgrounds. Some studies employ DNA molecular graph representations to capture structural information about the DNA, allowing for more nuanced analysis **[8]**. The choice of model architecture like Convolutional Neural Networks (CNNs) are well-suited for analyzing spatial hierarchies in data as CNNs can capture local patterns in DNA sequences. Implementation of bi-modal CNN and Long short-term memory (LSTM) could also achieve better classification results by learning intricate DNA structural and sequence features **[9]**. In addition to this, Zheng et. al., 2023 **[10]** used attention mechanism with CNN and LSTM for better classification performance **[10]**. Transformers **[11]**, which has gained prominence in various domains, offers advantages in capturing long-range dependencies in sequential data **[12-14]**. Bidirectional Encoder Representations from Transformers (BERT) **[15]** and its adaptations leverage the transformer model to enhance context understanding in sequence data **[15]**. Yet, most ML/deep learning (DL) approaches for DNA methylation site discovery fail to provide condition-specific methylation, focusing only on the likelihood of methylation for given sequences, regardless of condition. They overlook the influence of gene expression dynamics, which play a key role in determining differential methylation **[16-21]**. Existing tools rely heavily on sequence data without incorporating gene expression information. Some of the recent works on plant methylation modeling using machine learning are MGF6mARice **[8]**, DeepPGD **[9]**, MaskDNA-PGD **[10]**, Deep6mAPred **[12]**, i6mA-Vote **[22]**, iDNA-ABF **[23]**, MuLan-Methyl **[24]**,, MethBERT **[25]**, CpG-transformer **[26]**, DeepSignal **[27]**, PlantDeepMeth **[28]**, and MethNet **[29]**. But all of them suffer from the same limitations. To address this gap, integrated approaches should be developed which combine both DNA sequence and transcriptome data to capture the dynamic nature of methylation processes.

Based on the above concept, we have developed an innovative approach called **D**NA **M**ethylation **R**ecognition **U**nit **(DMRU)**. This system integrates a concurrent bi-modal architecture that combines DenseNet and Transformer encoder-decoder system within a generative deep learning framework. It has utilized 258 WGBS-seq and 326 RNA sequencing (RNA-Seq) data specifically from the plant species *Arabidopsis thaliana* and *Oryza sativa*. The primary objective of DMRU is to understand and learn from the interdependent variability between the methylation states present in the potential promoter regions of the genes (2kb upstream) and their corresponding RNA expression levels under specific environmental conditions. This enables DMRU to identify the most promising cytosine methylation points in the given sequence, independent of any species-specific models. In essence, DMRU’s design allows it to generalize across different plant species, providing a versatile tool for researchers in plant epigenetics.

The results achieved with DMRU is groundbreaking. In a rigorous benchmarking study, the system has consistently performed with an accuracy always exceeding 90%, and in cross-species universal manner. One of the distinguishing features of DMRU is its commitment to explainability in machine learning. By employing Gradient Weighted Class Activation Mapping (Grad-CAM) **[30]**, DMRU also provides insights into the underlying most important factors responsible for the observed methylation pattern for any given specific region. In contrast to existing software solutions, DMRU represents a pioneering effort in the realm of deep learning for plant epigenetics, while being the first auto-generative deep learning system specifically designed to decode condition specific cytosine methylation for any gene, with complete independence from any species specific model. Also, DMRU emerges as a promising alternative to costly and time consuming bisulfite sequencing experiments. **Figure 1** illustrates an abstract view of the introduction of the problem taken here and the solution.

## Materials and Methods

### Dataset retrieval, processing, and construction

For the development of universal and generalized model for the identification of methylated cytosines, we retrieved RNA-seq and WGBS-seq data for *Arabidopsis thaliana* and *Oryza sativa* from NCBI Sequence Read Archive (SRA). A total of 326 RNA-seq fastq data were collected from SRA databases for *A. thaliana* and *O. sativa*. Genomic sequences, annotations, and reference DNA sequences were downloaded from Ensembl Plants. Trimmomatic v0.39 **[31]** and in house developed reads processing tool, filteR **[32]**, were used to filter out poor quality reads, read trimming, and for adapter removal. Filtered reads were mapped back to the genome using Hisat2 **[33]**, afterwards read counting was done by using Rsubread **[34]**. Furthermore, the read counts were normalized to fragments per kilo base per million mapped reads (FPKM) for paired end reads and reads per kilo base per million mapped reads (RPKM) for single end reads.

In a manner similar to the RNA-seq, we collected a total of 258 WGBS-seq fastq data from SRA databases for *A. thaliana* and *O. sativa*. It has to be noted that, only those WGBS-seq fastq files were downloaded whose corresponding RNA-seq fastq files were available with the same experimental condition. Trimmomatic v0.39 **[31]** and filteR **[32]** were used to filter out poor quality reads, read trimming, and for adapter removal. The filtered reads were subsequently aligned to the reference genome using Bismark v0.22.2 **[35]**, retaining only uniquely aligned reads and discarding ambiguous mappings. For methylation analysis, we utilized Samtools v0.1.9 **[36]** to sort reads by genomic coordinates and remove PCR duplicates from the Bismark output, and ultimately converting this processed sequence alignment map (SAM) file to binary alignment map (BAM) output. The output BAM files were processed for methylation extraction using Bismark, excluding low-quality reads, and methylation sites (CG, CHH, CHG) were identified. Promoter sequences were extracted from gene transfer format (GTF) files, with methylated cytosines marked with a distinct character **“M”**, and corresponding gene expression profiles were integrated into the dataset. The dataset having sequences from *A. thaliana* has been named as Dataset “A” and the dataset having sequences from *O. sativa* has been named as Dataset “B” in the current study. Both these datasets were split in a 70:30 ratio for training and testing purposes, including for 10-folds random train:test trials. All details on data processing are illustrated in **Figure 2**. Complete list of every single samples (fastq files) which includes various sources, data volume and read count, experimental conditions, number of samples condition wise, are available in **Supplementary Table S2 Sheet 1-4**. Additionally, details of methylation site distribution (methylated and non-methylated sites) in all the three contexts (CG, CHH, and CHG) are also provided in this supplementary table.

**Figure 2:**
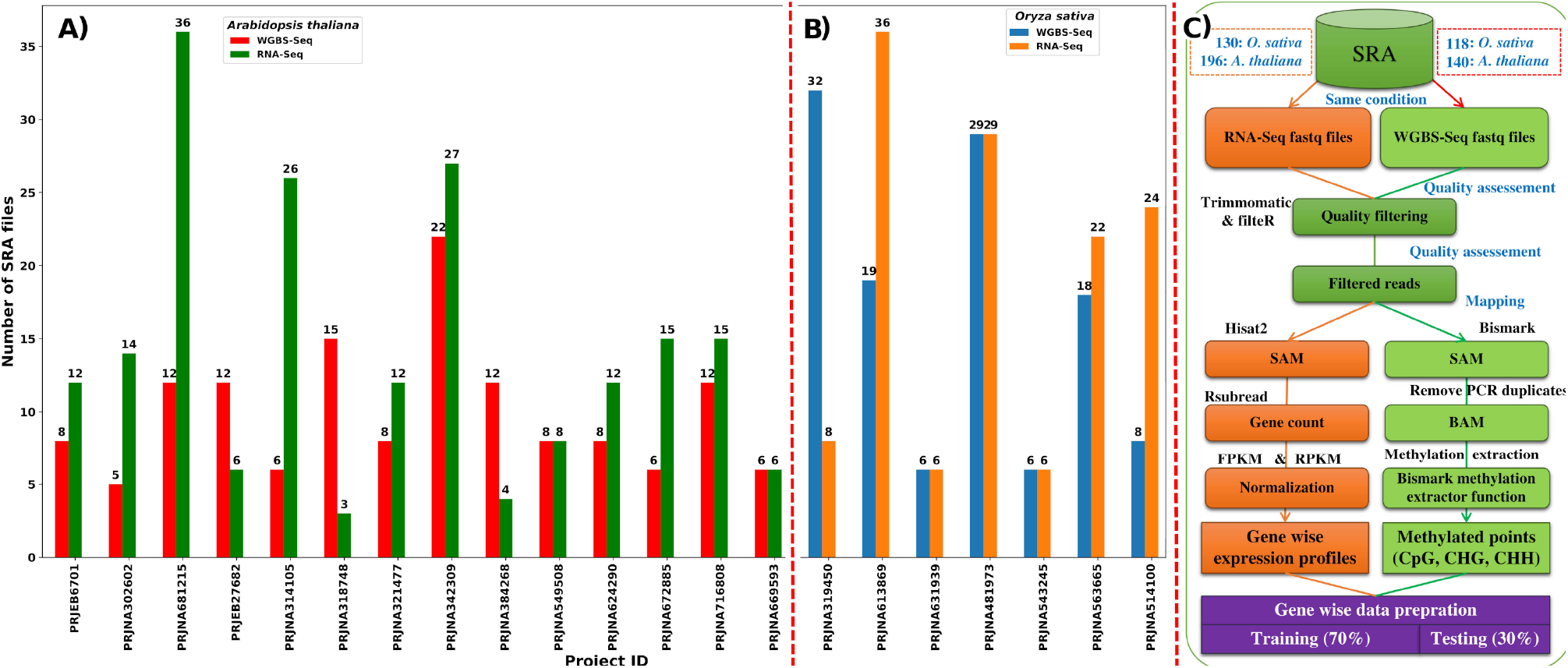
Data sources and processing. **a-b)** Distribution of WGBS and RNA-sequencing studies retrieved for the study collected from different projects from NCBI. **c)** Flowchart representation of dataset formation. The Dataset “A” and “B” were created for two plant species, *Arabidopsis thaliana* and *Oryza sativa*. Both the datasets contained the instances originating from RNA-seq and WGBS-seq, respectively.

### The Transformer-DenseNet system

We extracted gene expression data from RNA-seq for various conditions. This dataset encompasses the expression levels of each gene across different experimental contexts, providing a rich source of information to analyze how expressions of the target gene as well as other genes also correlate with methylation states. To process this data, we developed a DenseNet **[37]** model combined with a Transformer encoder to integrate gene expression and DNA methylation data for capturing local and global gene patterns.

### Constructing the DenseNet architecture to capture and connect the expression data

The DenseNet **[37]** model processes a 38,912-element tensor representing gene expression profiles, with each element corresponding to a gene’s expression level. Every such location is unique to a specific gene, when a user provide the transcriptome data, based on homology these gene specific cell are activated with their corresponding expression profiles. The DenseNet consists of convolution layers, batch normalization, max-pooling, four dense blocks, and three transition layers, totaling 121 layers, where dense blocks facilitate feature reuse and transition layers reduce dimensionality. In a dense block, each layer *“****l***” receives direct input from all preceding layers, producing output through operations like convolution and normalization.

Expressed mathematically, let “*H*_*l*_*”* represent the output feature maps of layer “*I*”. The output of layer “*I*” is computed as:

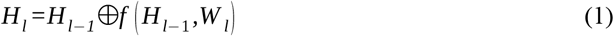

Here, “⨁ signifies concatenation, “*W*_*l*_*”* denotes the layer’s weights, and “***f***” incorporates batch normalization (BN), rectified linear units (ReLU), and convolution operations.

The growth rate, denoted by the hyperparameter “*k*” in DenseNet, emerges as a pivotal determinant in the architecture’s extraordinary performance. At each layer, “*k*” feature maps contribute to this global state, where the total number of input feature maps*”F*_*m*_*”* at the “*l*^*th*^*”* layer is calculated as:

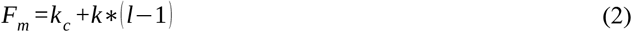

The term *“k*_*c*_*”* denotes the total number of channels in the input layer. DenseNet consists of dense blocks, where several convolutional layers are stacked together, followed by transition layers that reduce the dimensionality and down-sample the feature maps. This architecture’s output is intertwined through a bi-modal architecture in a feed-forward layer within the transformers encoder to capture RNA expression relationship with input target sequence data.

### Word representations of sequence data for Transformer

Each input sequence can be viewed as a collection of independent words. The DNA alphabet consists of four nucleotides: A, T, G, and C. By pairing these nucleotides with an additional character **“M”** for the methylated cytosines allows creating 3,125 unique 5-mer words, 15,625 unique 6-mer words, and 78,125 unique 7-mer words. In processing these sequences, overlapping windows of pentamers, hexamers, and heptamers could capture critical information about structural and functional regions within the DNA **[13-14, 38-42]**. Transformers takes two inputs i.e., source and target input. The source input consists of promoter sequences without any information of methylated cytosines with a maximum length of 5,985 words and 2,000 bases. The target input involves promoter sequences, which include methylation information with maximum length of 1,994 words and also spans 2,000 bases. To process these sequences, each unique word is assigned a distinct integer token. Once tokenized, these sequences are transformed into numeric vectors and matrices through a process known as embedding. By feeding these embedded sequences into the model, the Transformer can learn to translate sequences effectively.

### Implementation of the Transformers Encoders-Decoders

The encoder component processes the input sequence by transforming it into a series of continuous representations. It does this through several layers, each consisting of multiple attention heads and feed-forward neural (FFN) networks. The decoder, on the other hand, generates the output sequence based on the encoded information. It uses the context derived from the encoder’s outputs and incorporates its own attention mechanisms to focus on relevant parts of the input sequence while generating each token of the output.

### The Encoder

The encoder uses a multi-headed attention mechanism with self-attention layers to focus on relationships between words in the input, capturing long-range dependencies and contextual connections. The input sequence of word tokens is denoted as *“****Z****”*, with a length of *“****I****”*. The embedding layer projects these discrete input tokens into continuous vector embeddings. If “***v****”* is the size of the vocabulary and “***d****”* is the dimensionality of the word embeddings, the embedding layer is represented as a matrix *E*_*embed*_ ∈ℝ^*v*×*d*^ .

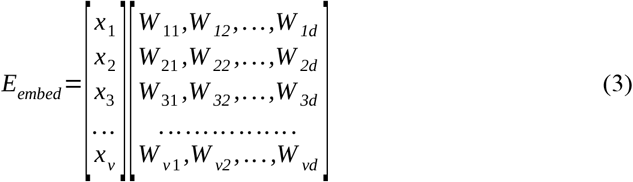

Each word in matrix “*E*_*embed*_*”* combines with its positional embedding “*P”*, where “*P”* shares dimension “*d”* with the word embedding vector. The resulting matrix, “*E*’_*embed*_*”*, is obtained through “*E’*_*embed*_ *=E*_*embed*_ *+P* “, where “*P”* is calculated using sinusoidal positional embedding equations:

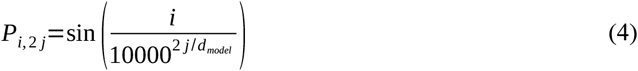

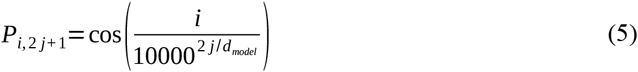

Here, “*i*” represents the position of the token, and “*d*_*model*_” is the embedding dimension. “*E*’_*embed*_” matrix enters the transformer encoder block, which processes it through a multi-head attention layer. This module involves multiple heads, each dividing its query, key, and value parameters N-ways and independently processing the splits. The output from each head produces an attention score derived from the key, query, and value computations. The model generates attention vectors through self-attention to capture relationships within the data, which are concatenated and passed through a feed-forward network with dropout layers to prevent overfitting. After normalization, the output integrates with the DenseNet output, containing gene expression data, and is fed into the transformer’s decoder block.

### Decoder block of the transformer

The decoder block in the transformer model facilitates sequence generation by using masked multi-head self-attention, attention over the encoder’s output, and position-wise feed-forward networks, with residual connections and layer normalization. This structure allows the decoder to align generated tokens with input representations, ensuring contextually relevant output.

### Masked Multi-Head Attention of Decoders

The primary purpose of masking in a decoder is to prevent the model from accessing future tokens when predicting the next token in a sequence. For each distinctive head “***i***”, symbolizing *i=* 1, 2, . .. ., *h*, the calculation of three pivotal matrices ensues: “*Q*_*i*_” (Query), “*K*_*i*_” (Key), and “*V*_*i*_” (Value). This is accomplished through the matrix multiplication of “*E*” with the corresponding weight matrices “*W*_*qi*_”, “*W*_*ki*_”, and “*W*_*vi*_”. The formulation of attention scores “*Z*_*i*_” transpires through the application of the softmax function to the scaled dot-product attention mechanism, adhering to the expression:

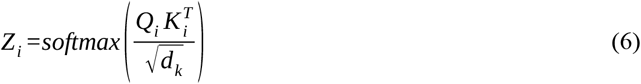

To preserve causality during training, a masking mechanism is applied to the attention scores, specifically designed to impede the model from attending to future positions, encapsulating the essence of masked self-attention. The ensuing phase involves the computation of the weighted sum, where each head “*i*” contributes to the formation of “*Head*_*i*_” via the formula:

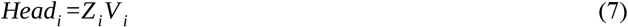

A concatenation of the outputs from all heads happens, culminating in a subsequent multiplication with the output weight matrix “*W*_*o*_”, yielding the definitive output of the Masked Multi-Head Attention mechanism in the decoder:

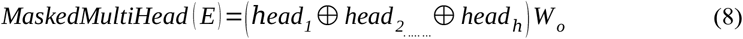

This exposition details the steps involved in using masked self-attention across multiple heads within the decoder of a Transformer model. During the decoding phase, the Masked Multi-Head Attention ensures future tokens are hidden, followed by integrating the encoder’s output for further processing. The Decoder Multi-Head Attention then focuses on relevant input elements, followed by feed-forward and normalization layers, the final output is evaluated to estimate conditional probabilities using softmax function, guiding the selection of contextually relevant tokens for sequence generation. The conditional probability *P* ⟨ *y*_*t*_|*y*_1_, .…,*y*_*t* −*1*_, *x* ⟩ for the next token “*y*_*t*_” is computed using the following equation:

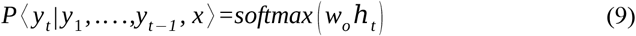

where, “*w*_*o*_” is the weight matrix, and “*h*_*t*_” is the hidden state at position “*t*”. This process repeats iteratively for each time step until the entire sequence is decoded. The model uses the generated tokens as input for subsequent time steps, allowing for autoregressive decoding.

Developed by Papineni et al, in 2002 **[43]**, Bilingual Evaluation Understudy (BLEU) has become a standard benchmark for assessing the quality of translations by comparing them to reference translations. The final BLEU score is computed as the weighted geometric mean of the precision scores of 5-grams levels in the generated sequence compared to the reference sequence, adjusted by the brevity penalty as given below:

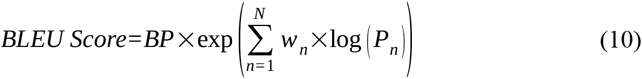

where, “*BP*” is the Brevity Penalty, accounting for the length of the generated sequence relative to the reference. “*N*” is the maximum order of 5-grams considered. “*w*_*n*_” is the weight assigned to each n-gram precision. “*P*_*n*_” is the precision of 5-grams in the predicted sequence. This formula quantifies the precision of the model’s output against reference sequences, considering 5-gram orders and assigning appropriate weights. **Figure 3** depicts the operation of the implemented deep-learning encoder-decoder system.

**Figure 3:**
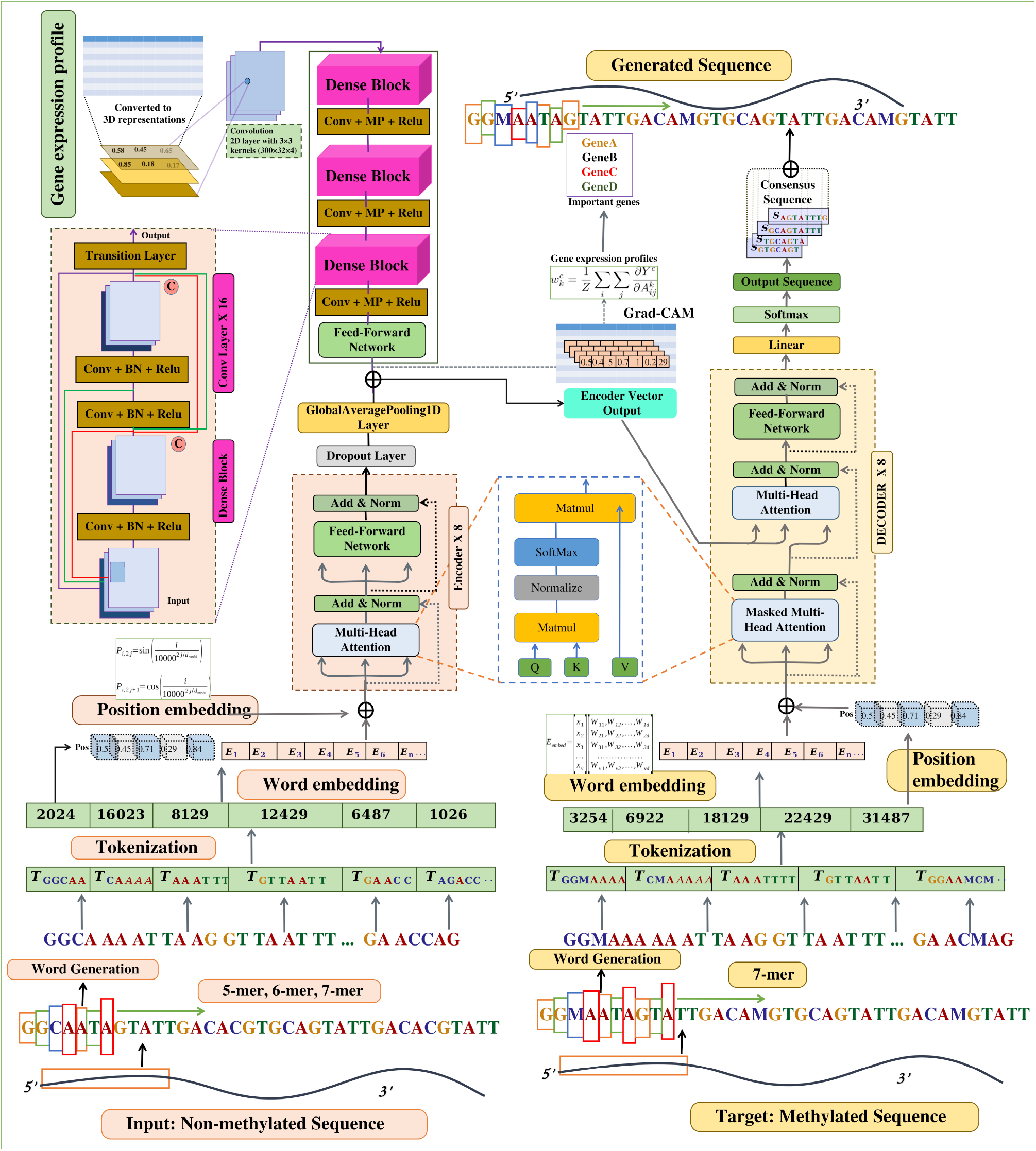
Implementation of the DMRU Deep Co-learning system using Transformers and DenseNet to annotate DNA methylation. The encoder processes the promoter sequences, without methylation representations, in penta, hexa, and heptamer words representations, while the decoder receives the methylated sequences representations for the corresponding sequence. Eight layers of encoder:decoder pairs were incorporated. In the parallel, a DenseNet consisting of 121 layers accepts input in the form of the associated RNA expression profile. The learned representations from the encoder and DenseNet are merged and passed to the encoder-decoder multi-head attention layer, which also receives input from decoder layers. Subsequently, the resulting output undergoes conditional probability estimation, playing a pivotal role in the decoding process. Following layer normalization, the model proceeds to calculate the conditional probability distribution over the vocabulary for the next token. Finally, the resultant tokens were converted back into words, representing the methylated states for the given sequence.

### Evaluation of the generated output from the Transformer-Decoder

The decoder of the transformer generates output sequences representing both methylated and unmethylated cytosines, which are then matched to target sequences. Post-decoding, the model’s accuracy is evaluated by comparing the generated sequences to experimental data on cytosine methylation, analyzing base-by-base agreement. The model was trained using 70% of Dataset “A” and tested on the remaining 30%, with performance further validated through 10-fold random independent trials. Overfitting was assessed using mean absolute error (MAE) values, ensuring consistency and robustness across variable datasets.

Full method details are provided in the **Supplementary File 1 Materials**. The readers are highly encouraged to consult the **Supplementary File 1** for in depth details of methods and data.

## Results and Discussion

### Data collection and formation of datasets

DNA methylation has been reported to be a function of sequence properties and several genes **[1, 16-21, 44-49]**. Some of well identified such genes in plants are: DNA methyltransferase MET1, DDM1, VIM1–3, chromomethylase CMT1-3, histone methyltransferases kryptonite (KYP), Su(var)3-9 homologue 5 (SUVH5), IDN2, IDN2, DRM2, and SUVH6 **[16-21]**. However, these are usually fundamental genes which are not condition specific but usually expressed in a uniform way. While the fact is that in plants the regulatory DNA methylations are highly spatio-temporal, suggesting involvement of several other molecular factors in making the epigenetic regulation highly specific and time controlled.

In this study, we have investigated the relationship between DNA methylation profiles and transcriptome expression profiles through two important plant species, *Arabidopsis thaliana* and *Oryza sativa*. Our investigation was centered around the 2-kilobase (kb) upstream regions of every gene as the potential promoter regions where DNA methylations at cytosines have been reported critical in regulating the downstream gene’s expression. Each promoter sequence was represented multiple times, because of varying methylation states of cytosines across different conditions. In terms of scale, the dataset for *O. sativa* comprised 55,42,110 promoter sequences with variable methylation states, covering 38,756 genes and 40 conditions while *A. thaliana* contributed 51,14,144 promoter sequences with variable methylation states, covering 32,368 genes and 45 conditions. These data also used the corresponding gene expression data for the given condition from condition specific transcriptome profile. The average coverage of the RNA-seq data was ∼17 for *A. thaliana* and ∼8 for rice. These datasets have been named as Dataset “A” (*A. thaliana*) and Dataset “B” (*O. sativa*) in the current study, also are illustrated in **Figure 2** and details of the data provided in **Supplementary Table 2 Sheet 1-4**. A total of 45 experimental condition for *Arabidopsis* and 40 experimental condition for rice were taken.

### Conjoint deep-learning on sequence properties and conditional transcriptome profiles raises an efficient model to detect cytosine methylation

The implementation of the DMRU model represents a significant advancement, particularly in the study of gene regulation through the interplay of DNA methylation, methylation region’s sequence information, and surrounding transcriptome’s profile including the expression of the downstream gene. Central to this implementation is a composite deep co-learning system combining Transformers and DenseNet architectures to capture complex patterns in gene promoter methylation and RNA expression variability under different environmental conditions. The efficacy of the model was evaluated using Dataset “A”, with a 70:30 training-testing split, ensuring no overlap or redundancy between the sets. The training phase involved 1,200 distinct promoter sequences paired with RNA-seq data across 45 experimental conditions. After processing through the integrated Transformers-DenseNet system, it achieved an impressive 90.31% accuracy **(Figure 4a)** on an unseen test set of 200 promoter sequences across 45 experimental conditions covering a total of 31,600 different methylation profile sequences, demonstrating its ability to identify methylated cytosines using RNA-seq data **(Figure 4a)**. The model’s performance was further improved through hyperparameter optimization, which is detailed in the following section and **Supplementary Table 2 Sheet 5**.

**Figure 4:**
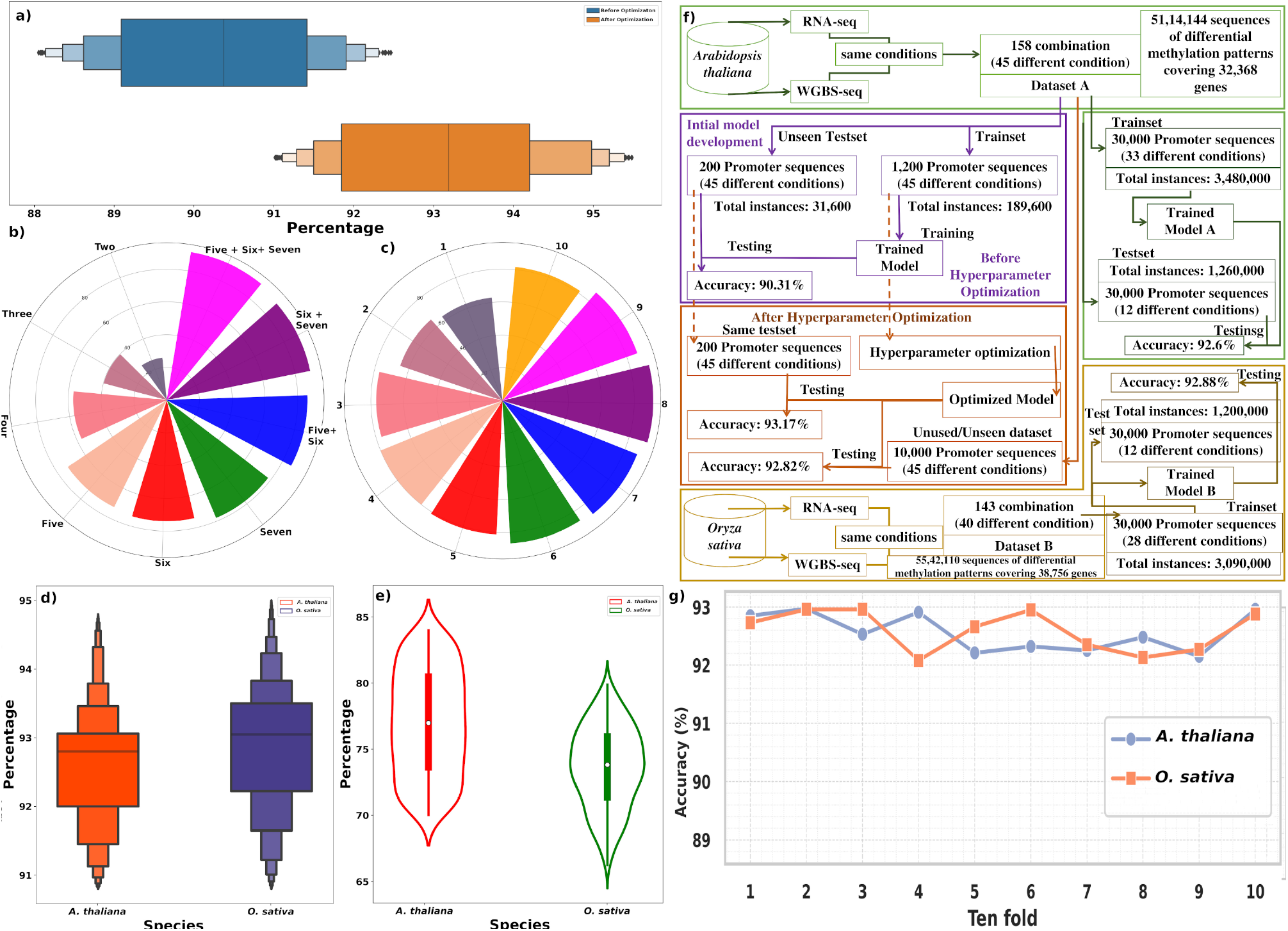
Comprehensive performance assessment of the raised deep co-learning models. **a)** The model was evaluated on an unseen test set consisting of randomly selected 200 promoter sequences from Dataset “A” before and after hyper-parameter optimization. **b)** Ablation analysis for six main properties was conducted to evaluate their impact on model performance. These words representations appeared highly additive and complementary to each other as the performance increased substantially as they were combined together. **c)** Hyper-parameters optimization effect on the performance due to varying numbers of encoders and decoders for the Transformers-DenseNet system. At the addition of the eighth encoder-decoder pair, a significant change in the model’s ability to handle the complexities was observed. **d-e)** Self and cross-species performance of DMRU across plant species *A. thaliana* and *O. sativa*. For this evaluation, we utilized two distinct datasets, referred to as Dataset “A” and Dataset “B”, comprising sequences from two model plant species, *A. thaliana* and *O. sativa*. The approach involved training a model using sequences from *A. thaliana* and subsequently testing it on the sequences from *A. thaliana and O. sativa*, and vice-versa. This self and cross-species assessment were critical to ascertain the accuracy and adaptability of the models across different genomic contexts. **f)** Flow diagram depicting the various stages in the model development and corresponding data distribution. **g)** Ten fold random independent train:test trails of Model “A” and “B”.

To identify the key features contributing to the model’s accuracy in decoding cytosine methylation, an ablation analysis was performed, evaluating the impact of different sequence-based features as mentioned in the methods section. The hypothesis was that integrating these representations would enhance the model’s performance by providing more contextual information. Results showed that integrating higher-order words improved performance significantly **(Figure 4b)**. Dimers achieved 26.02% accuracy, trimers 39.8%, pentamers reached 72.77%, hexamers 74%, and heptamers 78%. However, computational complexity and memory usage increased with longer representations, as seen when training with heptamers, which required 46 GB of GPU memory and 1,456 seconds per epoch. Combining different representations further boosted accuracy, with the pairing of hexamers with heptamers reaching 89.32%. The best result, 91.48% accuracy, was achieved by combining pentamers, hexamers, and heptamers. This combination demonstrated the additive benefits of information sharing, highlighting the importance of integrating multiple representations for optimal model performance. The findings are illustrated in performance plots **(Figure 4b)** and detailed in **Supplementary Table 2 Sheet 6**, which clearly show the improvements in accuracy associated with various combinations of sequence representations.

### Hyper-paramter optimization further improved the model

Hyperparameter optimization was crucial to enhancing the performance of the deep-learning model, which initially used values derived from previous work **[13-14, 38, 50]**. The first step involved determining if the identified sequence and expression properties formed the right features, followed by finalizing the deep-learning architecture. Once a satisfactory baseline accuracy was achieved, hyperparameter optimization was applied to improve the performance. Exploring different numbers of encoder-decoder pairs with different numbers of attention heads in the Transformer-DenseNet system revealed that performance peaked at the eighth encoder-decoder pair, after which additional layers led to diminishing returns **(Figure 4c)**. Same number of attention heads was found performing best. Dropout layers (with fractions of 0.12 and 0.14) were used to regularize the model and reduce overfitting, while normalization ensured stability across layers. The architecture incorporated a feed-forward network with 48 and 64 nodes with Relu and Selu activation function, respectively, enhancing feature extraction and transformation. The output from DenseNet was processed through the Transformer encoder to capture variability in RNA expression and methylation data. In training, a masked multi-head attention mechanism prevented information leakage, and the decoder’s feed-forward layer generated coherent outputs. The model was optimized using the Adam optimizer (learning rate of 0.001), trained for 100 epochs with a batch size of 4 (**Figure 3)**. To prevent overfitting, we applied weight decay with a coefficient of 1e-5. This helped improve the performance of the model. We also implemented early stopping with a patience of 10 epochs, monitoring validation loss using PyTorch function ReduceLRonPlateau. Training was halted if the validation loss did not improve for 10 consecutive epochs, a standard practice, which helped prevent overfitting and reduce the training time. After hyperparameter tuning, the model achieved an accuracy of 93.17%, a 3% improvement over its baseline performance **(Figure 4a and Supplementary Table 2 Sheet 5). Supplementary Table 2 Sheet 7** provides a list of all key hyperparameter implemented during summarization while **Supplementary Table 2 Sheet 8-10** and **Supplementary Figure S1** provides the training curves (loss/accuracy/gpu usage vs. epochs) during training and testing process. In addition to it, training this Transformer-DenseNet on 2,00,000 sequences with an NVIDIA RTX A5000 GPU reached 84–96% utilization and upto 22.7 GB of memory. The AMD EPYC 7302 CPU peaked at 68% due to parallel data preprocessing, while memory usage hit 61.4 GB from caching and tensor handling. In contrast, inference/test was far less demanding with only 3.2 GB GPU memory, below 20% GPU use, and just 18% CPU usage making DMRU feasible to run on even low configuration local systems. However, DMRU is implemented as a web-server, where all these matters don’t matter.

### Remarkably consistent performance observed across wide range of experimental data

By evaluating upon 10,000 promoter sequences from Dataset “A” that were not included in the training and testing phases, the model achieved an impressive accuracy of 92.82% **(Figure 4d)**. Following this promising result, we developed two co-learning models, Model “A” from Dataset “A” and Model “B” from Dataset “B”. We further tested these models on the testset of Dataset “A” and Dataset “B”, each comprising a total of 30,000 unique promoter sequences corresponding to *A. thaliana* and *O. sativa*, respectively. Both the model’s performance remained consistent, with an average accuracy of 92.74% across the datasets. More specifically, the accuracy for *A. thaliana* was recorded at 92.6% from Model “A”, while for *O. sativa* from Model “B”, it slightly improved to 92.88% **(Figure 4d). Figure 4f** provides the details of data splitting, subsequent stages in model development and noted performance leap. For detailed gene and condition wise metrics focusing on per-cytosine accuracy (includes accuracy, precision, recall, and F1-score alongside confusion matrices) specifically quantify how well the model translates unmethylated input sequences into correctly methylated output sequences, **Supplementary Table 3 Sheet 1** is provided. Afterwards, we assessed contribution from the integration of the RNA expression profile handled through DenseNet, the DL system was raised again without it. This analysis marked decrease in the model’s accuracy, plummeting to 67.23% (p-value >> 0.01). Such a significant decline suggested for the necessity of jointly analyzing sequence data and RNA expression profiles which ultimately enriched the model’s performance. The 10-fold random trials of training and testing showed consistent performance for Dataset “A”, with negligible MAE difference between training and testing sets (0.0029) (**Supplementary Table S2 Sheet 11)**. A t-test confirmed no significant overfitting. For all these 10 cross-validations, the model consistently performed above 92% accuracy, suggesting a highly stable and consistent performance. **Figure 4g** provides the result of 10 fold random trails of train:test for both species. Through out the study the data were ensured to avoid any overlap or redundancy, ensuring unbiased performance testing.

### Accurate cross-species identification of DNA methylation where GC similarity is the key, not homology

Detecting DNA methylation computationally itself is a big challenge, whose next level comes with condition specific detection of DNA methylation. It becomes top challenge when one looks for developing a tool which could work cross-species in total universal manner also. Unfortunately, most of these challenges have remained unaddressed, which DMRU has successfully solved now.

The DMRU model, initially species-specific, was tested for cross-species applicability by evaluating its performance on sequences from *A. thaliana* and *O. sativa* using the two datasets. When models trained on one species were tested on the other, the accuracy was lower than within-species performance, with the *O. sativa* model achieving a maximum accuracy of 83.9% on *A. thaliana* sequences, and the *A. thaliana* model reaching 79.89% on *O. sativa* sequences **(Figure 4e)**. To improve cross-species performance, the initial approach relied on sequence homology, but this only achieved around 83% accuracy. Both results clearly highlights that the raised models left enough scope for further improvement as they were far below than the accuracy levels observed for within species runs.

To develop a universal model capable of identifying DNA methylation in the promoter sequences from gene expression data across various species, our initial approach centered on leveraging homology aimed to harness evolutionary relationship between the sequences where the query sequences/genes were looked for their corresponding homologous ones from *A. thaliana*. However, rigorous testing, the model underperformed, achieving only ∼83% accuracy **(Figure 5a-b)**. This outcome prompted us to rethink our approach as well as highlighted how wrong one can go while traditionally relying on the homology, especially for epigenetic controls. In epigenetic control it is GC isochore which should be looked for instead of homology as it determines the methylation rate and pattern **[51-53]**. Therefore, the GC content (GC%) of the 2 Kb upstream regions for the genes were evaluated in both *A. thaliana and O. sativa*. By grouping genes with similar GC% value ranges, we created a new model based on these clusters. This innovative approach proved fruitful when we applied the GC%-based model to the datasets from both species, as the results significantly improved. The model derived from *A. thaliana* demonstrated an impressive accuracy exceeding 90% on *O. sativa* (Dataset “B”), while the model based on *O. sativa* also achieved above 90% accuracy on *A. thaliana* (Dataset “A”) **(Figure 5a-b)**. These outcomes underscore the critical role which GC content plays in the context of gene regulation and methylation.

**Figure 5:**
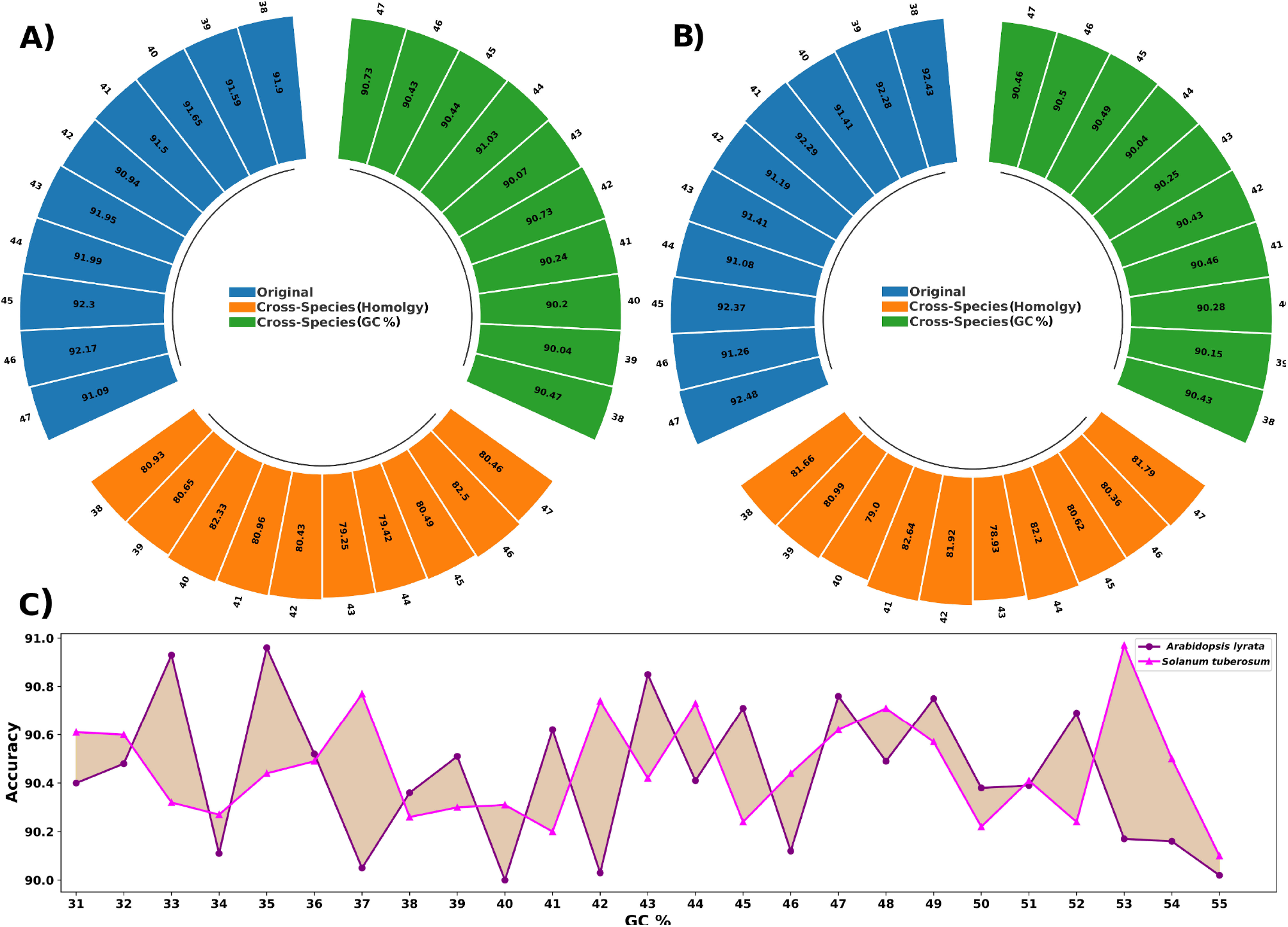
Performance comparisons for the GC% based universal model of DMRU across species. Performance assessment for the model raised using **a)** *Arabidopsis thaliana* data, and **b)** *Oryza sativa* data, with self genes (Blue coded), homology based cross-species (orange coded), and GC% similarity based cross-species testing. **c)** Cross-species performance of DMRU based on GC % clusters. Here, GC% cluster based models from *A. thaliana* was tested across *A. lyrata* and *S. tuberosum* genes. Consistently >90% accuracy was observed for every GC% clusters. The GC% of the 2 Kb upstream regions for the genes were evaluated in both *A. thaliana and O. sativa*. By grouping genes with similar GC% value ranges, we created new models based on these clusters, the results were significantly improved. These results underscore the critical role which GC content plays in the context of gene regulation and methylation.

In the next step, the above analysis was extended forward to identify cross-species DNA methylation on other unseen species. Based on experimental data availability for validation, this was done for *Arabidopsis lyrata* and *Solanum tuberosum*, for which WGBS-seq and RNA-seq data were available. Here also, the DMRU model developed from Dataset “A” achieved an average accuracy >90% for both the species, affirming its efficacy in handling cross-species conditions. **Figure 5c** provides the comparison details for both species by DMRU for cross-species performance. All these series of validation studies reinforced that DMRU performed consistently with high accuracy and emerged as a promising first of its kind tool for cross-species methylation identification without any dependency on predefined models while implementing generative and explainable AI innovative way.

### Why GC% similarity worked? And what could be the biological implications?

GC content in the first gate of entry for the transcriptional regulation **[54-55]**, the pioneer factors, which read the methylation pattern, DNA shape, and GC contour to open the DNA for migrant and settler TFs which are DNA motif dependent unlike the pioneers. GC content directly influence these patterns **[51-52]**.

Further to this, evolutionary relationship between GC content and methylation pattern has been established **[53]**. Besides these direct implications GC content drift also affects the availability of TF binding sites. An analysis with the various GC content categories taken in this study, it was found that genes belonging to same GC content category displayed strongest expression correlation for the consisdered 45 conditons in *Arabidopsis*. This strong correlation vanished gradually as one drifted aparts to more different GC content (**Supplementary Figure S2**).

All this strongly reason, why GC content similarity worked to derive a universal cross-species model to detect differential methylation pattern.

### DMRU outstands in comparative benchmarking studies

To evaluate the performance of DMRU, we compared it with nine different tools: DeepPGD **[9]**, MaskDNA-PGD **[10]**, iDNA-ABF **[23]**, MuLan-Methyl **[24]**, DeepSignal **[27]**, PlantDeepMeth **[28]**, MethNet **[29]**, iDNA-MS **[56]**, and iDNA-ABT **[57]**. The four out of the compared nine tools viz. iDNA-MS, DeepPGD, MuLan-Methyl, and MethNet had bugs to run **(Supplementary Table S4 Sheet 1)**. All these five tools were tested across datasets “A” and “B”. The performance on the test sets from datasets “A” and “B”, containing 12 different conditions, showed that DMRU outperformed all compared algorithms across various performance metrics. Unlike the other tools, which provide uniform methylation profiles across all conditions, DMRU excels at identifying condition-specific DNA methylation, making it more applicable in real-world scenarios. None of the existing tools are capable for condition specific DNA methylation prediction. As highlighted in **Figure 6a-c and Supplementary Figure S3**, all these tools output exactly the same result and fail to capture the dynamic condition specific methylation patterns in *O. sativa*. On the other hand **(Figure 6d)**, DMRU captured the conditions specific methlyations in these regions. **Supplementary Figure S3-4** also presents dynamic condition specific methylation patterns in *A. thaliana*. **Supplementary Figure S5-7** show that the compared tools did not breach even 80% accuracy mark when the accuracy was calculated for every sequence under different conditions.

**Figure 6:**
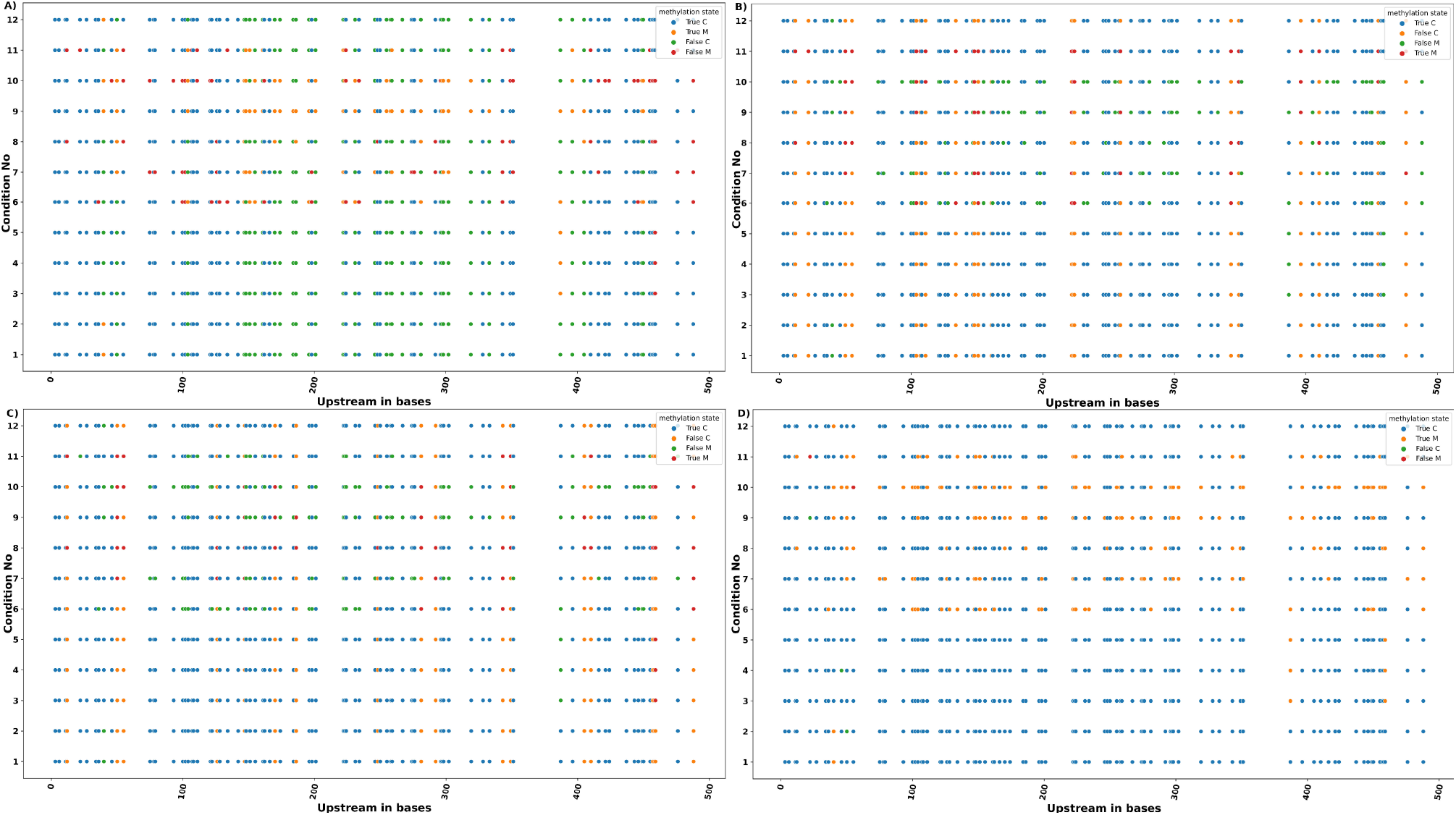
Performance benchmarking of the universal model of DMRU based on GC% for *Oryza sativa*. The performance over the 12 different experimental conditions for the compared software tools were benchmarked for the upstream region (have 500 bases shown) of gene Os01g0110100, for the tools **(a)** iDNA-ABF, **(b)** iDNA-ABT, **(c)** MaskDNA-PGD, and **(d)** DMRU. Here also, comparative benchmarking returned outstanding results, reinforcing DMRU as the most reliable software too to detect differential methylation. T-test based p-values for comparison between software tools, DMRU vs iDNA-ABT: 1.49e-05, DMRU vs iDNA-ABF: 1.14e-05, and DMRU vs MaskDNA-PGD: 1.97e-05. As can be seen, other tools keep giving same methylation patterns in all experimental conditions, while DMRU very accurately captured the condition specific methylation patterns.

This also needs to be noted that cross-species performance of DMRU stood-out, as showed above for the *A. thaliana* raised model accurately detecting the differential methylation in a monocot, *Oryza sativa*. Yet, to further verify it, the DMRU with *A. thaliana* raised model was run across *Z. mays* genome, a distant monocot species, for three different experimental conditions and their associated methylome **(Supplementary Table S4, Sheet 3)**. Here also DMRU retained the same performance level, and breached 90% accuracy, clearly reinforcing its universal capabilities. Also, it maintained the same level of lead from the compared tools, none of which could even cross-beyond 80% accuracy (**Supplementary Figure S8**).

With this all, DMRU stands at present the only approach which could capably suggest differential methylation patterns across different conditions, emerging also as a good alternative for costly bisulfite sequencing experiments.

### Explainable deep learning gives insight into the most critical condition specific factors functioning for DNA methylation

DMRU has implemented Grad-CAM **[30]** scoring scheme to pull out the genes from the transcriptome profile which appear to be critical for condition specific methylation for any given sequence stretch. This approach involves computing the gradient of a target class, denoted as “***c***” alongside the global average of the activation feature map “***F***” derived from the first convolutional layer “***l***”. By synthesizing these gradients with the activation maps, DMRU generates class-discriminative profiles which highlight the importance of specific features, expressing genes from the transcriptome profile in the present condition. The mathematical formulation and operational principles of Grad-CAM are described as follows:

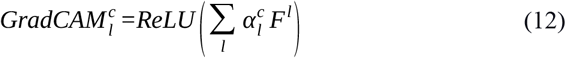

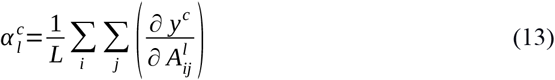

Where, 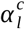 denotes the neuron importance weights, highlighting the most relevant features that contribute to the models accuracy. **‘*L’*** is the vector length in the feature map, and 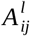 represents the activation at **“*ij***^***th***^**”** position in the input array. Full details of Grad-CAM implementation is covered in **Supplementary File 1 Materials and Methods** section.

To illustrate the effectiveness of the implemented approach, case of gene AT1G13450 from *Arabidopsis thaliana* has been considered as example here. This gene is pivotal due to its role in binding to one of the cis-acting elements, BoxII, which resides within the upstream promoter region of light-responsive genes, regulating plant growth. In this context, we utilized Grad-CAM to visualize the influence of gene expression on DNA methylation. To extract relevant features, we employed the first convolutional layer of our Transformer-DenseNet model, effectively enabling Grad-CAM to elucidate crucial gene features linked to DNA methylation. The findings of this analysis is illustrated by **Figure 7a** where the top 100 genes ranked by their Grad-CAM scores across multiple conditions were clustered and presented. To further contextualize our findings within biological frameworks, we conducted Gene Ontology (GO) analysis on the genes identified through Grad-CAM. GO analysis of these high Grad-CAM scoring top 100 genes unveiled significant insights that a large proportion of these genes were intricately linked to essential plant developmental processes **(Figure 7b-d)**.

**Figure 7:**
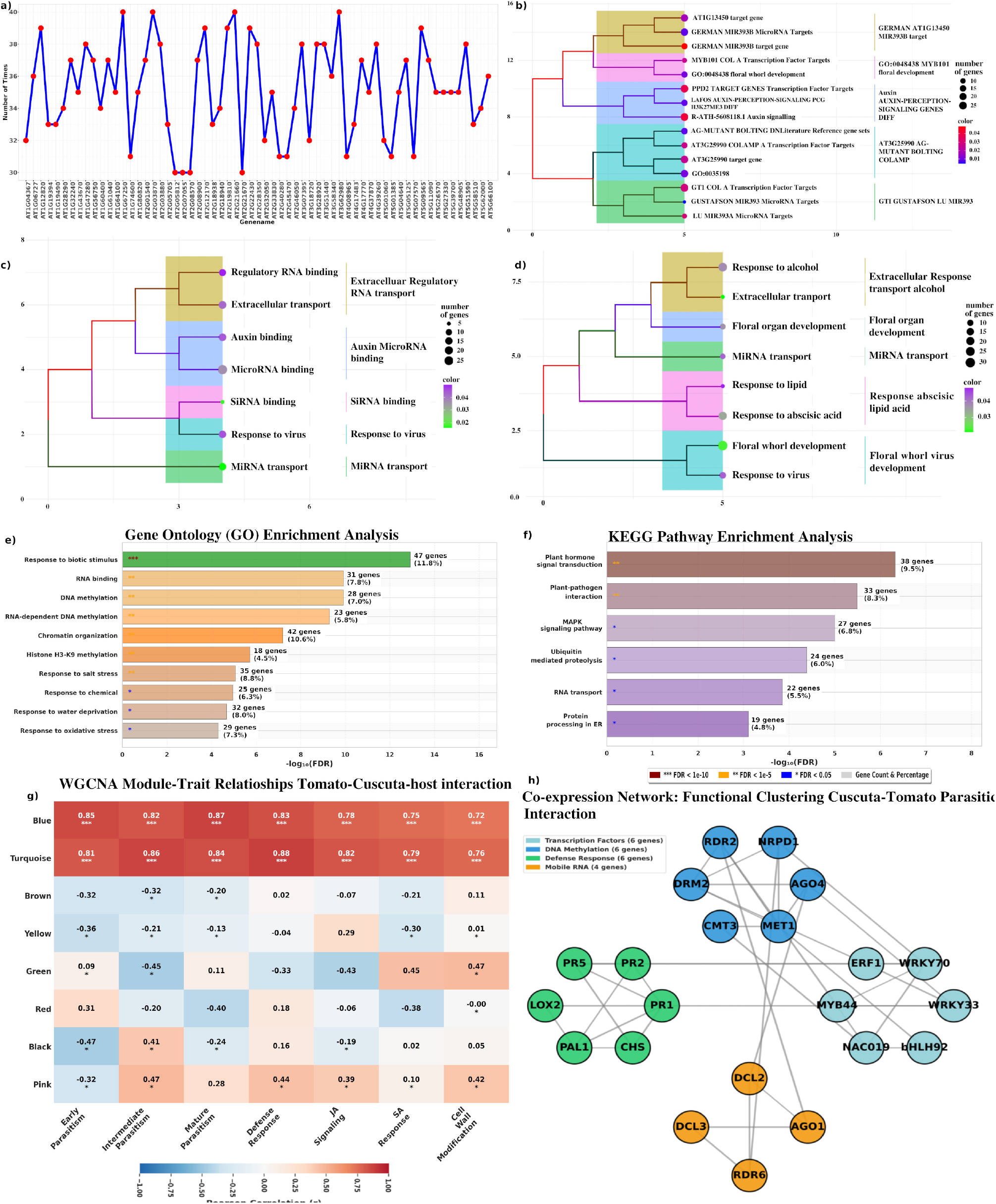
Explainability of DMRU system. It helps to understand the most impactful genes features possibly responsible for the observed methylation patterns. **(a)** The top 100 genes ranked by their Grad-CAM scores for their methylation impact across multiple conditions were clustered for the target gene AT1G13450, **(b-d)** Gene Ontology (GO) analysis on the genes identified through Grad-CAM. This application demonstration while considering this target gene aptly proposed the relevant genes which concur very well with respect to their common functions. Conditions specific differential methylation appears to control specific functionality. **(e-h) Functional and network characterization of Grad-CAM–prioritized, DMRU-identified promoters that overlap WGCNA stress modules in the Cuscuta–tomato interface. (e)** GO enrichment for the gene set defined by the intersection of DMRU/Grad-CAM high-scoring promoters (threshold ≥4) and stress-correlated WGCNA modules. Bars show enrichment strength as −log10(FDR); numbers to the right of bars report gene counts and percentage of the overlapping set. GO terms are ordered by significance and highlight processes tightly linked to epigenetic regulation and stress responses. Significance symbols (legend) indicate FDR thresholds used to mark highly significant terms (*** FDR < 1×10^−10^; ** FDR < 1×10^5^; * FDR < 0.05). **(f)** KEGG pathway enrichment for the same overlapping gene set. Bars represent −log10(FDR) and annotated gene counts. **(g)** WGCNA module–trait heatmap showing Pearson correlations between module eigengenes and parasitism/stress traits sampled across haustorial developmental stages. Correlation coefficients are annotated in each cell (color scale: blue = negative, red = positive). The two modules most strongly associated with parasitic stress (labelled Blue and Turquoise) display consistently high positive correlations across interface stages (representative r ≈ 0.72–0.87), supporting their designation as stress-responsive modules. **(h)** Co-expression network extracted from the stress-associated WGCNA modules, functionally clustered and annotated to emphasize core epigenetic players and defense components. Node color denotes functional class; node labels display gene symbols for key loci. Edge connections represent strong weighted co-expression/topological overlap, illustrating putative regulatory links among methylation enzymes, RdDM factors, histone modifiers, transcription factors.

Notably, they played vital roles in maintaining RNA binding, facilitating floral whorl developmental mechanisms, and driving stamen development, which is crucial for reproduction in flowering plants **(Figure 7c)**. This emphasizes the foundational importance of these genes in not just growth, but also in the adaptations of *A. thaliana*. In *Arabidopsis*, floral organs develop in concentric whorls, and AT1G13450 acts as a regulator of genes that determine the identity of these organs. Specifically, it affect the expression of genes responsible for establishing the correct patterning of the second and third whorls. It also plays role in controlling floral symmetry, the proper arrangement of floral organs and it also helps in establishing floral symmetry, an essential feature of floral development **[58-59]**. AT1G13450 is involved in interactions with different TFs such as LEAFY (LFY) and APETALA1 (AP1), which are known to regulate floral organ identity **[59]**. The interaction of these transcription factors can influence the development of floral organs in *Arabidopsis* too. AT1G13450 contributes to the integration of various signals needed for the correct positioning and formation of floral organs. Moreover, our analysis revealed that these genes exhibit enrichment in various molecular functions. Key activities identified include extra-cellular transport, floral whorl and organ development, response to abscisic acid, which are essential for signaling pathways that regulate growth and development, indicative of their roles in biochemical reactions crucial for metabolism and synthesis **(Figure 7c-d)**. The examination of KEGG pathways further corroborated our findings, illustrating the involvement of these genes in these important metabolic pathways, as depicted in **Figure 7b-d**.

This application demonstration while considering gene AT1G13450 clearly displayed the level of accuracy and importance DMRU is capable to attain, while churning out relevant information also. The Grad-CAM part of DMRU very aptly proposed the relevant genes in the system which could be involved with this gene where DNA methylation appears to have significant stake in its regulation.

### Biological interpretation and utility of Grad-CAM-derived influential genes: A case study on Cuscuta-Tomato parasitic stress

In this case study, publicly available RNA-seq data from Jhu et al., 2022 **[60]** (PRJNA687611, PRJNA756681) which capture transcriptome dynamics at the *Solanum lycopersicum*–*Cuscuta campestris* interface across early, intermediate and mature haustorial stages, were used to investigate parasitism-associated regulatory programs. These datasets represent stress-induced conditions where Cuscuta parasitism triggers host defense responses, including epigenetic changes. We pre-processed the data, performed weighted gene co-expression network analysis (WGCNA) using the R package (v1.72-5) to identify stress-responsive modules. DMRU was applied to identify the condition-specific methylation patterns in the promoter regions (2 kb upstream) of tomato genes. Grad-CAM scores from DMRU were used to rank the influential genes, and mapped these to WGCNA modules for overlap analysis.

WGCNA identified two modules (blue and turquoise) highly correlated with parasitic stress traits (Pearson correlation >0.8, p < 0.001; total 437 genes), enriched in defense pathways like jasmonic acid (JA) signaling, salicylic acid (SA) responses, and cell wall modification (GO:0006952, defense response; KEGG:04626, plant-pathogen interaction) **(Supplementary Table S4 Sheet 4)**. DMRU’s Grad-CAM analysis on these data yielded high-scoring genes (threshold >4), with 91% (398 genes) of the reported stress modules overlapping with DMRU. This overlap was highly significant (hypergeometric test: p = 1.23 × 10^-45^), indicating Grad-CAM effectively captured stress-relevant regulators. GO and KEGG enrichment of the overlapping genes highlighted epigenetic regulation associated e.g., RNA binding (GO:0003723, p=3.4 × 10^-12^), response to biotic stimulus (GO:0009607, p=1.2 × 10^-15^), and phenylpropanoid biosynthesis (KEGG:00940, p=4.7 × 10^-8^), which are linked to methylation and histone modifications under stress (e.g., homologs of DRM2, AGO4) **(Figure 7e-h)**. Interestingly, in their study the authors had experimentally validated 12 genes and implicated them in Cuscuta induced stress, causing DNA methylation. All of these 12 genes were detected by DMRU Grad-CAM as influential genes associated with the observed DNA methyaltion patterns. All this reinforced the great credibility and utility of DMRU and its Grad-CAM to detect condition specific DNA methylation and infer associated influencer genes. This all also opens a plethora of opportunities and experimental interactions. Further details on this study is given in **Supplementary File 1**.

The DMRU system is freely accessible at https://scbb.ihbt.res.in/DMRU/, and **Figure 8** provides an overview of DMRU webserver implementation. Full details on the server part are provided in the **Supplementary File 1 Results**.

**Figure 8:**
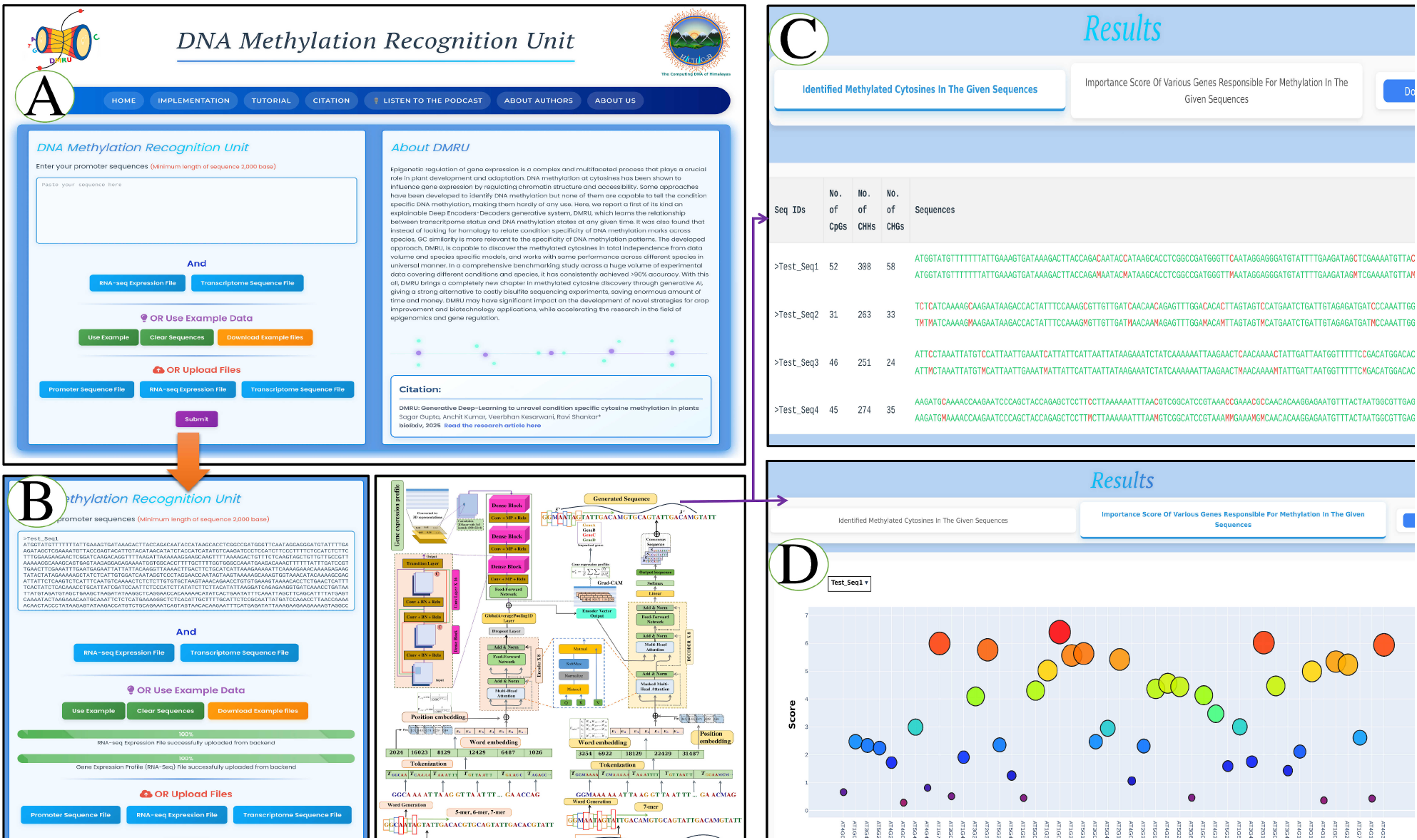
DMRU webserver implementation. **a)** Input data box where the user can either paste or load the input files, **b)** Inputs are DNA sequence in FASTA format and gene expression profile in “tsv” format output from RNA-seq data analysis. **c)** The methylated cytosines are represented in an interactive form. Download option for the result in the tabular format. **d)** Also, important feature scoring is implemented and is represented in the form of interactive bubble plot depicting the distribution of scoring distribution for the most influential genes associated with the observed DNA methylation pattern for the user provided target sequence.

## Conclusion

DNA methylation at cytosines regulate the downstream gene to attain spatio-temporal gene expression pattern. This is very crucial for development, survival, and adaptations mechanism of plants. Currently, only though Bisulfite based sequencing experiments one can find the methylation of DNA and its differential patterns, making its detection a costly affair. No existing software tools are capable to tell the differential methylation patterns. Here, we have develpoed a deep-learning based encoder:decoder system in generative manner to suggest the condition specific cytosine methylation with consistently high accuracy. It was tested across a large number of experimental data, covering several conditions. It also worked with same level of accuracy in cross-species manner. DMRU now makes it possible for anyone to find out the methylation pattern of their target genes promoter for any given condition. One just needs to load the RNA-seq data of the condition and the target sequences. DMRU goes further and also tells about the most influential genes associated with the observed methylation patterns. DMRU results can minimize the experimental loads. One can carry out precise guided experiments like CRISPR based editing at influenced methylation, carry out methylation sensitive essays in targetted manner with DMRU result, map its results with chromatin accessibility data, and several other experiments. DMRU opens a completely new sets of doors in plant regulomics studies, as well as epigenetic studies in total. With this all, DMRU emerges as a first of its kind software tool, and a much cheaper, faster, and accurate alternative to costly and time consuming experiments like Bisulfite-seq.

## Declarations

### Software and Data availability

All the secondary data used in the present study were publicly available and their due references and sources have been provided in **Supplementary Table S1-4**. The software has also been made available as a webserver at https://scbb.ihbt.res.in/DMRU. The pseudocode of the implementation has also been provided as **Supplementary File 1** under section: “**Pseudocode of the Transformer-DenseNet model architecture**”.

### CRediT authorship contribution statement

**Sagar Gupta:** Data curation, Software, Methodology, Validation, Visualization, Resources, Formal analysis, Writing – original draft. **Anchit Kumar:** Data curation, Visualization, Formal analysis. **Veerbhan Kesarwani:** Data curation, Formal analysis. **Umesh Bhati:** Formal analysis. **Ravi Shankar:** Conceptualization, Formal analysis, Investigation, Methodology, Project administration, Supervision, Writing – original draft, Writing – review & editing.

## Acknowledgements

SG, AK, and VK are thankful to DBT, India for financial support as project associateship. UB is thankful for DBT SRF fellowship. SG, VK, and UB are also thankful to Academy of Scientific and Innovative Research (AcSIR) for their Ph.D. enrollment. All authors are thankful to the Director, CSIR-IHBT, for his kind support for this study. This MS has CSIR-IHBT MSID **5833**.

## Key Points

1. Existing tools rely heavily on sequence data without incorporating gene expression information. And, this is why most of them fail to predict differential methylation in different conditions.
2. For the first time here, to capture the dynamic nature of methylation processes both DNA sequence and transcriptome data has been learned through a Transformer-DenseNet Deep-Learning system. This has enabled the methylation studies in plants free from the above mentioned bottlenecks and makes it feasible to detect methylated cytosines with accuracy and reliability for plant genome.
3. The developed tool, DMRU, has been tested across a huge volume of experimental data where it breached the accuracy of 93% and always scored >90% in every validation test,, even cross-species.
4. DMRU is expected to revolutionize the plant regulatory research as it will empower to mimic costly high-throughput experiments like WGBS-seq with its highly accurate methylated cytosine identification for any given condition for plant genome. This work can also be extended to animal systems.

## Author Biographies

Sagar Gupta is a PhD research scholar in Bioinformatics at Studio of Computational Biology & Bioinformatics, HiCHiCoB, CSIR-Institute of Himalayan Bioresource Technology (CSIR-IHBT), India. His research interests are focused on computational regulomics with machine and deep learning.

Anchit Kumar is a research scholar in Bioinformatics at Studio of Computational Biology & Bioinformatics, HiCHiCoB, CSIR-Institute of Himalayan Bioresource Technology (CSIR-IHBT), India. His research interests are focused on machine and deep learning.

Veerbhan Kesarwani is a PhD research scholar in Bioinformatics at Studio of Computational Biology & Bioinformatics, HiCHiCoB, CSIR-Institute of Himalayan Bioresource Technology (CSIR-IHBT), India. His research interests are focused on structural and molecular dynamics aspects in regulation.

Umesh Bhati is a PhD research scholar in Bioinformatics at Studio of Computational Biology & Bioinformatics, HiCHiCoB, CSIR-Institute of Himalayan Bioresource Technology (CSIR-IHBT), India. His research interests are focused on regulatory systems biology.

Ravi Shankar is a senior principal scientist and Coordinator of the Himalayan Centre for High-throughput Computational Biology (HiCHiCoB), a National Bioinformatics Center (BIC) of Department of Biotechnology, Ministry of Science & Technology, at Division of Biotechnology, CSIR-Institute of Himalayan Bioresource Technology (CSIR-IHBT), India. He has more than 20 years of research experience in Computational Genomics, Regulomics, and Machine Learning, where he has successfully supervised 10 PhDs towards completion, developed more than 20 software tools, and received more than 10 research grants as PI. He has been conferred with several prestigious recognition including NIH, USA project fellowship and INSA & Australian Academy of Science Fellowship. More about his research group can be found at https://scbb.ihbt.res.in.

## Supplementary Information

**Supplementary Figure S1: Training and validation loss, accuracy, and GPU usage over epochs. a)** This line plot illustrates the loss progression during training and validation phases across epochs. The training loss is shown in solid red, and the validation loss is shown in dashed green. The loss values are tracked to evaluate model convergence and potential overfitting. **b)** This figure shows the training and validation accuracy (%) across epochs. Accuracy values are displayed as percentages, reflecting the model’s classification performance during each training epoch. **c)** This plot depicts GPU utilization (%) during training and validation over time.

**Supplementary Figure S2: Two randomly selected genes from GC% group of 42% GC content were initially selected (G3:AT5G18755, G4:AT4G00520)**. Within group, we observed high expression correlation between the member genes. These two genes too displayed the same (r ≈ 0.85). When these genes were compared with other GC% groups, the expression similarity gradually vanished as the GC dissimilarity increase. In the present figure we randomly selected two representative genes for each GC% group.

**Supplementary Figure S3: The performance over the 12 different experimental conditions for the compared software tools were benchmarked for the upstream region (have 500 bases shown) of gene AT1G01010 and Os01g0110100, for the tools (a, c) DeepSignal, and (b, d) PlantDeepMeth**.

**Supplementary Figure S4: Performance and benchmarking of the universal model of DMRU based on GC% for *Arabidopsis thaliana***. The performance over the 12 different experimental conditions for the compared software tools were benchmarked for the upstream region (have 500 bases shown) of gene AT1G01010, for the tools **(A)** iDNA-ABF, **(B)** iDNA-ABT, **(C)** MaskDNA-PGD, and **(D)** DMRU.

**Supplementary Figure S5: Performance accuracy of the benchmarked software tools (DMRU and iDNA-ABT) across different conditions**. iDNA-ABT did not breach even 80% accuracy mark while DMRU easily crosses 90% when the accuracy was calculated for every sequence under different conditions. Rendering DMRU effective across different experimental conditions.

**Supplementary Figure S6: Performance accuracy of the software tools across different conditions. (A)** iDNA-ABF and **(B)** MaskDNA-PGD. These software tools did not breach even 80% accuracy mark when the accuracy was calculated for every sequence under different conditions.

**Supplementary Figure S7: Performance accuracy of the software tools across different conditions. (A)** DeepSignal and **(B)** PlantDeepMeth. These software tools did not breach even 80% accuracy mark when the accuracy was calculated for every sequence under different conditions.

**Supplementary Figure S8: Performance accuracy of the software tools across different conditions on *Zea mays***. Software tools other than DMRU did not breach even 80% accuracy mark when the accuracy was calculated for every sequence under different conditions.

**Supplementary Figure S9: Comparison of GC content and methylation levels across transcription factors binding sites (TFBSs)**. The line plot shows the GC percentage distribution (GC%) of TFBS sequences in *Arabidopsis thaliana*. The scatter plot displays the corresponding methylation percentage for each TF (in red). A green regression line indicates the overall pattern of methylation changes with increasing GC%. Higher GC is mostly associated with higher chance of DNA methylation.

**Supplementary Figure S10: Comparative analysis of transcription factor (TF) binding and expression correlation in genes grouped by GC content in *Arabidopsis thaliana* and *Oryza sativa***. The left side represents genes from *A. thaliana* belonging to the 35% GC content group. Promoter regions (2 kb upstream) of these genes were analyzed for TF binding sites, revealing that AT3G24120 and BIM2 together cover over 80% of the gene set. Expression correlation (Pearson’s r) between TFs and selected genes within the group is shown in the matrix, with strong positive correlations observed, indicating strong co-regulation. The right side shows the analysis in *O. sativa*, where the homologous gene (OsNAC06) belongs to the 56% GC content group. Promoter analysis of this group did not identify any binding sites for the homologous TFs. Corresponding expression correlation between these TFs and selected target genes including (OsNAC06) shows no strong correlation, suggesting a loss of regulatory interaction likely due to the GC content shift across the compared species.

**Supplementary Table S1: Brief list of some of the published tools for DNA methylation identification**.

**Supplementary Table S2: Information regarding to (A)** dataset “A” and “B” general information, **(B)** Hyperparameter optimization and details of Transformers, **(C)** Details of abylation and tenfold MAE analysis, **(D)** sRNA-seq, **(D)** Statistics of the training curves (loss/accuracy/gpu usage vs. epochs) during training and testing process.

**Supplementary Table S3: Statistics of the key metrics such as AUC, precision, recall, or F1-score are shown across all test conditions**.

**Supplementary Table S4: Statistics of the bugs identified during benchmarking and data regarding GC% and methylation content in transcription factor binding sites**.

