## Supplementary File 1 for "DMRU: Generative Deep-Learning to unravel condition specific cytosine methylation in plants"

[1] Studio of Computational Biology & Bioinformatics,
The Himalayan Centre for High-throughput Computational Biology,
(HiCHiCoB, A BIC supported by DBT, India), Biotechnology Division,
CSIR-Institute of Himalayan Bioresource Technology (CSIR-IHBT),
Palampur (HP), 176061, India.

[2]Academy of Scientific and Innovative Research (AcSIR),
Ghaziabad, Uttar Pradesh- 201002

### Materials and Methods

#### Dataset retrieval, processing and construction

For the development of universal and generalized model for the identification of methylated cytosines, we retrieved RNA-seq and WGBS-seq data for *Arabidopsis thaliana and Oryza sativa* from NCBI Sequence Read Archive (SRA). A total of 326 RNA-seq fastq data were collected from SRA databases which comprises 196 and 130 different read files for *A. thaliana* and *O. sativa*, respectively (**Supplementary Table S2 Sheet 1-2**). A total of 45 and 40 different experimental conditions were retrieved for *A. thaliana* and *O. sativa*, respectively. Genomic sequences, annotations and reference DNA sequences were downloaded from Ensembl Plants. Trimmomatic v0.39 **[1]** and in house developed reads processing tool, filteR **[2]**, were used to filter out poor quality reads, read trimming, and for adapter removal. Filtered reads were mapped back to the genome using Hisat2 **[3]**. After mapping, read counting was done by using Rsubread **[4]**. Furthermore, the read counts were normalized to fragments per kilo base per million mapped reads (FPKM) for paired end reads and reads per kilo base per million mapped reads (RPKM) for single end reads, using in-house developed shell scripts for every single gene across different conditions.

In a manner similar to the RNA-seq, we collected a total of 258 WGBS-seq fastq data from SRA databases having 140 and 118 different read files for *A. thaliana* and *O. sativa*, respectively (**Supplementary Table S2 Sheet 3-4**). It has to be noted that, only those WGBS-seq fastq files were downloaded whose corresponding RNA-seq fastq files were available with the same experimental condition. Trimmomatic v0.39 **[1]** and filteR **[2]** were used to filter out poor quality reads, read trimming, and for adapter removal. The filtered reads were subsequently aligned to the reference genome using Bismark v0.22.2 **[5]**, retaining only uniquely aligned reads and discarding ambiguous mappings. For methylation analysis, we utilized Samtools v0.1.9 **[6]** to sort reads by genomic coordinates and remove PCR duplicates from the Bismark output, and ultimately converting this processed SAM file to BAM output. Afterwards, the output BAM files underwent methylation extraction using the Bismark methylation extractor function, with reads exhibiting a conversion rate below 90% or fewer than three non-converted cytosines in non-CpG contexts excluded from analysis. Furthermore, we employed the CX report file generated by Bismark, which provide details of the three types of methylation sites (CG, CHH, CHG). We extracted the two kilobases (kb) upstream promoter sequences for each gene of both the species from the downloaded gene transfer file (GTF) file. Additionally, using this CX report file, we marked all methylated cytosines within the promoter sequences with a distinct character **"M"** to differentiate the methylated cytosines with unmethylated cytosines. For every promoter single sequences, corresponding gene expression profiles were also considered in the development of this dataset. The dataset having sequences from *A. thaliana* has been named as Dataset "A" and the dataset having sequences from *O. sativa* has been named as Dataset "B" in the current study. Both these datasets were split in a 70:30 ratio for training and testing purposes, including for 10-folds random train:test trials. All details on data processing are illustrated in **Figure 2**. Complete list of every single samples (fastq files) which includes various sources, data volume and read count, experimental conditions, number of samples condition wise, are available in **Supplementary Table S2 Sheet 1-4**. Additionally, details of methylation site distribution (methylated and non-methylated sites) in all the three contexts (CpG, CHH, and CHG) are also provided in this supplementary table.

#### The Transformer-DenseNet system

We extracted gene expression data from RNA sequencing (RNA-seq) for various conditions. This dataset encompasses the expression levels of each gene across different experimental contexts, providing a rich source of information to analyze how expressions of the target gene as well as other genes also correlate with methylation states. To process this data, we developed a DenseNet **[7]** model architecture known for its efficiency in handling vanishing gradient and retention of learnings from previous layers. We established a bi-modal architecture with DenseNet and Transformer encoder which effectively combines expression profiles and DNA methylation in the target sequence region information. This architecture enables the model to take sequences and gene expression data as input, allowing it to capture both local and global patterns in gene expression that may be linked to methylation. The expression input to the DenseNet is a vector with a size of 38,912 elements corresponding to the positions of genes holding their respective expression values for the given condition. The DenseNet architecture we developed for this study was implemented using PyTorch.

#### Constructing the DenseNet architecture to capture and connect the expression data

Introduced by Gao Huang and colleagues in 2017, DenseNet **[7]** is a multi-layered CNN architecture that enhances feature propagation and reuse, addressing challenges like the vanishing gradient problem commonly encountered in deep networks. This dense connectivity allows gradients to flow more easily through the network during training, resulting in improved convergence and performance. The DenseNet model developed here processes a tensor of 38,912 elements representing gene expression profiles 38,912 genes (each element corresponds to the positions of genes, each associated with its respective expression level). Every such location is unique to a specific gene, when a user provide the transcriptome data, based on homology these gene specific cell are activated with their corresponding expression profiles. This architecture consisted a convolution layer with 32 convolution filters (kernel size = 3), batch normalization, and 2D max-pooling (stride = 2), and incorporates four dense blocks and three transition layers, totaling 121 layers. Dense blocks, fostering feature reuse through dense connectivity, play a crucial role. In a dense block, each layer "*l*" receives direct input from all preceding layers, producing output through operations like convolution and normalization. Transition layers effectively reduce the dimensions before the next dense block.

Expressed mathematically, let "$H_l$" represent the output feature maps of layer "*l*". The output of layer "*l*" is computed as:

$$H_l = H_{l-1} \oplus f\left(H_{l-1}, W_l\right) \tag{1}$$

Here, " $\oplus$ " signifies concatenation, "$W_l$" denotes the layer's weights, and "***f***" incorporates batch normalization (BN), rectified linear units (ReLU), and convolution operations. The unique dense connectivity enhances feature reuse, contributing to the model's ability to learn complex hierarchical features effectively.

The growth rate, denoted by the hyperparameter "*k*" in DenseNet, emerges as a pivotal determinant in the architecture's extraordinary performance. DenseNet's innovative design treats feature maps as a global network state, enabling impressive results even with a smaller growth rate. At each layer, "*k*" feature maps contribute to this global state, where the total number of input feature maps "$F_m$" at the "$l^{th}$" layer is calculated as:

$$F_m = k_c + k * (l - 1) \tag{2}$$

The term "$k_c$" denotes the total number of channels in the input layer. DenseNet consists of dense blocks, where several convolutional layers are stacked together, followed by transition layers that reduce the dimensionality and down-sample the feature maps. These transition layers incorporate batch normalization, ReLU activation, and 1x1 convolutions, making the network both efficient and manageable. To enhance computational efficiency, a 1x1 convolution layer precedes each 3x3 convolution layer. This design, known as a bottleneck layer, reduces the number of input feature maps, typically exceeding the output feature maps "*k*", by generating 4k feature maps. This architecture's output is intertwined through a bi-modal architecture in a feed-forward layer within the transformers encoder to capture RNA expression relationship with input target sequence data to raise a bi-modal learning model to identify methylated cytosines in global and cross-species manner.

#### Word representations of sequence data for Transformer

The Transformer encoder-decoder architecture is particularly powerful due to its ability to process input and output sequences of varying lengths in a unified, end-to-end manner. This flexibility makes it well-suited for tasks that involve translating or transforming sequences, such as sequences transformations or natural language processing (NLP). Each input sequence can be viewed as a collection of independent words or tokens. The DNA alphabet consists of four nucleotides: A, T, G, and C. By pairing these nucleotides, we can create a total of 16 unique dimers. An alphabet of the four bases with an additional character **"M"** for the methylated cytosines allows creating 3,125 unique 5-mer words, 15,625 unique 6-mer words, and 78,125 unique 7-mer words. This exponential growth in words allows for a rich representation of genetic information. In processing these sequences, overlapping windows of pentamers, hexamers, and heptamers can capture critical information about structural and functional regions within the DNA **[8-14]**. Transformers takes two inputs i.e., source and target input. The source input consists of promoter sequences without any information of methylated cytosines with a maximum length of 5,985 words and 2,000 bases, providing a detailed representation of the DNA being analyzed. The target input typically involves promoter sequences, which include methylation information. This input has a maximum length of 1,994 words and also spans 2,000 bases. To process these sequences, each unique word is assigned a distinct integer token. This tokenization step is crucial as it converts the raw sequences into a structured format suitable for Transformers models. Once tokenized, these sequences are transformed into numeric vectors and matrices through a process known as embedding. The Transformer's encoder processes the source sequences while the decoder generates the target sequences based on the encoded information. By feeding these embedded sequences into the model, the Transformer can learn to translate sequences effectively. This setup allows for efficient handling of large datasets and complex sequence transformations.

#### Implementation of the Transformers Encoders-Decoders

The encoder component processes the input sequence by transforming it into a series of continuous representations. It does this through several layers, each consisting of multiple attention heads and feed-forward neural networks. The encoder is designed to capture hierarchical features, meaning it can identify patterns at different levels of abstraction. Its ability to consider the entire input sequence at once enables it to capture dependencies and relationships across the sequence. The decoder, on the other hand, generates the output sequence based on the encoded information. It uses the context derived from the encoder's outputs and incorporates its own attention mechanisms to focus on relevant parts of the input sequence while generating each token of the output.

#### The Encoder

The encoder utilizes a multi-headed attention mechanism that features self-attention layers by employing a dot-product method. These layers enable the encoder to focus on various words in the input, consider their relationships and how they relate to one another, when making decisions. This design allows the transformer model to effectively capture long-range dependencies and contextual connections within the input data. In the transformer architecture, the sequences derived from the tokenized representation of the previous step are input into the system's input layer. Each word is transformed into a word vector of a specified embedding size through the embedding process. This results in an output matrix that is structured by samples, sequence length, and embedding size, facilitating the model's ability to understand and manipulate the data.

The input sequence of word tokens is denoted as *"Z"*, with a length of *"I"*. The embedding layer projects these discrete input tokens into continuous vector embeddings. If "***v***" is the size of the vocabulary and "***d***" is the dimensionality of the word embeddings, the embedding layer is represented as a matrix $E_{embed} \in \mathbb{R}^{v \times d}$ .

$$E_{embed} = \begin{bmatrix} x_1 \\ x_2 \\ x_3 \\ \dots \\ x_v \end{bmatrix} \begin{bmatrix} W_{11}, W_{12}, \dots, W_{1d} \\ W_{21}, W_{22}, \dots, W_{2d} \\ W_{31}, W_{32}, \dots, W_{3d} \\ \dots\dots\dots\dots\dots \\ W_{v1}, W_{v2}, \dots, W_{vd} \end{bmatrix} \quad (3)$$

Each word in matrix "$E_{embed}$" combines with its positional embedding "*P*", where "*P*" shares dimension "*d*" with the word embedding vector. The resulting matrix, "$E'_{embed}$", is obtained through " $E'_{embed} = E_{embed} + P$ ", where "*P*" is calculated using sinusoidal positional embedding equations:

$$P_{i,2j} = \sin\left(\frac{i}{10000^{2j/d_{model}}}\right) \quad (4)$$

$$P_{i,2j+1} = \cos\left(\frac{i}{10000^{2j/d_{model}}}\right) \quad (5)$$

Here, "*i*" represents the position of the token, and "$d_{model}$" is the embedding dimension. "$E'_{embed}$" matrix enters the transformer encoder block, which processes it through a multi-head attention layer. This module involves multiple heads, each dividing its query, key, and value parameters N-ways and independently processing the splits. The output from each head produces an attention score derived from the key, query, and value computations.

Initially, the model generates individual attention vectors, which are essential for capturing relationships within the input data. This step involves applying self-attention to produce contextual embeddings for each token in the sequence. The attention vectors are then concatenated and subsequently forwarded to a feed-forward network block within the encoder. To combat the risk of overfitting, dropout layers were employed. Afterwards, a normalization layer implemented as Layer Normalization follows. This normalization helps stabilize the learning process by normalizing the outputs, ensuring that the mean and variance remain consistent throughout the training. The output post-normalization is then directed to fully connected feed-forward layers and a dropout layer is introduced to prevent overfitting, followed by another normalization layer. At this stage, the output is integrated with the DenseNet output, encapsulating vital gene expression information from the RNA-seq data. The final output from this enhanced encoder structure feeds into the decoder block of the transformer.

#### Decoder block of the transformer

The decoder block of the transformer architecture plays a crucial role in the overall functioning of the model, particularly in tasks involving sequence generation, such as text generation. Each decoder block consists of several key components: masked multi-head self-attention, multi-head attention over the encoder's output, and position-wise feed-forward networks, all combined with residual connections and layer normalization. The masked multi-head self-attention mechanism is essential for ensuring that the prediction for a particular token depends only on the tokens that have been generated so far, thus preventing any information leakage from future tokens. In contrast, the second attention layer in the decoder block attends to the encoder's output, enabling the decoder to incorporate information from the input sequence. This interaction allows the model to generate contextually relevant output by aligning the generated tokens with the input representations. The attention mechanisms in both layers operate on the principle of computing weighted averages of the input embeddings, where the weights are determined by the similarity between the query and key vectors, thus allowing the model to focus on different parts of the input and output sequences as needed.

**Masked Multi-Head Attention of Decoders**

The primary purpose of masking in a decoder is to prevent the model from accessing future tokens when predicting the next token in a sequence. This is essential for autoregressive tasks, where predictions for the current position should rely solely on previously generated tokens. By using a masking mechanism, the decoder adheres to this constraint, ensuring that it does not "cheat" by incorporating information from future positions. Masking is typically implemented using a triangular matrix, where positions that cannot be attended to are set to a large negative value. This allows the softmax function to produce zero probabilities for these positions, effectively ignoring them during the attention calculation. This strategic design promotes the autoregressive nature of decoding, ensuring the model generates sequences in a coherent manner.

For each distinctive head "$i$", symbolizing $i = 1, 2, \ldots, h,$ the calculation of three pivotal matrices ensues: "$Q_i$" (Query), "$K_i$" (Key), and "$V_i$" (Value). This is accomplished through the matrix multiplication of "$E$" with the corresponding weight matrices "$W_{qi}$", "$W_{ki}$", and "$W_{vi}$". The formulation of attention scores "$Z_i$" transpires through the application of the softmax function to the scaled dot-product attention mechanism, adhering to the expression:

$$Z_i = softmax\left(\frac{Q_i K_i^T}{\sqrt{d_k}}\right) \tag{6}$$

To preserve causality during training, a masking mechanism is applied to the attention scores, specifically designed to impede the model from attending to future positions, encapsulating the essence of masked self-attention. The ensuing phase involves the computation of the weighted sum, where each head "*i*" contributes to the formation of "$Head_i$" via the formula:

$$Head_i = Z_i V_i \tag{7}$$

A concatenation of the outputs from all heads happens, culminating in a subsequent multiplication with the output weight matrix "$W_o$", yielding the definitive output of the Masked Multi-Head Attention mechanism in the decoder:

$$MaskedMultiHead(E) = (head_1 \oplus head_2 \ldots\ldots \oplus head_h) W_o \tag{8}$$

This exposition details the steps involved in using masked self-attention across multiple heads within the decoder of a Transformer model. During the decoding phase, the process begins with the Masked Multi-Head Attention mechanism, which analyzes the input sequence while ensuring that future tokens remain concealed. Once this initial processing is complete, the output from the encoder is integrated as a secondary input for the decoder. Next, the Decoder Multi-Head Attention mechanism operates similarly to the encoder's attention, allowing the model to selectively focus on specific elements of both the input sequence and the encoder's output. Afterwards, the output is refined through two Feed-forward and normalization layer. Following these steps, the final output is evaluated to estimate conditional probabilities, which is a key part of the decoding process. Building on the normalized output, the model calculates a conditional probability distribution over the vocabulary for the next token in the sequence. This is accomplished using the softmax function, which converts the raw scores into a probability distribution. The resulting probabilities guide the selection of the next token, making this step essential for generating outputs that are coherent and contextually relevant. The conditional probability $P\langle y_t | y_1, \ldots, y_{t-1}, x \rangle$ for the next token "$y_t$" is computed using the following equation:

$$P\langle y_t | y_1, \ldots, y_{t-1}, x \rangle = softmax(w_o h_t) \tag{9}$$

where, "$w_o$" is the weight matrix, and "$h_t$" is the hidden state at position "$t$". This process repeats iteratively for each time step until the entire sequence is decoded. The model uses the generated tokens as input for subsequent time steps, allowing for autoregressive decoding.

Developed by Papineni et al, in 2002 **[15]**, Bilingual Evaluation Understudy (BLEU) has become a standard benchmark for assessing the quality of translations by comparing them to reference translations. The BLEU score quantifies how closely a machine-generated sequences output aligns with target sequences. It is a precision-based metric that primarily focuses on the overlap of n-grams (contiguous sequences of n items from a given sample of text) between the generated text and the reference texts. The score ranges from 0 to 1, where 0 indicates no overlap and 1 indicates perfect overlap.

The final BLEU score is computed as the weighted geometric mean of the precision scores of 5-grams levels in the generated sequence compared to the reference sequence, adjusted by the brevity penalty. Mathematically, this can be represented as:

$$BLEU\ Score = BP \times \exp\left(\sum_{n=1}^{N} w_n \times \log\left(P_n\right)\right) \tag{10}$$

where, "$BP$" is the Brevity Penalty, accounting for the length of the generated sequence relative to the reference. "$N$" is the maximum order of 5-grams considered. "$w_n$" is the weight assigned to each n-gram precision. "$P_n$" is the precision of 5-grams in the predicted sequence. This formula quantifies the precision of the model's output against reference sequences, considering 5-gram orders and assigning appropriate weights. The primary advantage of the BLEU score is its ability to provide a quick, quantitative assessment of translation quality. Its reliance on n-gram matching makes it easy to compute and interpret.

#### Evaluation of the generated output from the Transformer-Decoder

The decoder segment of the transformer is designed to generate multiple words as part of its output sequence comprising both the methylated and unmethylated cytosines bases for each provided sequence. The crucial step post-decoding involves matching these generated words with target sequences. To take the final decision on the full length promoter output sequence generated from the raised model, sequences were mapped back to the original target sequence and per base cytosines analysis was carried to compare the agreement with actual experimental data for methylated states of cytosines for the given experimental condition for every generated sequence. In this way, the accuracy of the raised model was calculated for the complete promoter output sequence. This entire part of the study has used 70% of the Dataset "A" as the training set to build the model and its performance on the remaining 30% totally unseen test dataset. **Figure 3** depicts the operation of the implemented deep-learning encoder-decoder system. Further to this, the evaluation was also done through 10-fold random independent train-test trial runs while building the model from the scratch every time maintaining the same 70:30 ratio of train:test datasets to evaluate the consistency of the developed method across variable learning and testing datasets.

#### Explainable deep learning gives insight into the most critical condition specific factors functioning for DNA methylation

Significant efforts are underway to enhance the transparency and interpretability of deep learning. DLs perform exceptionally good but are hard to explain the causality. Explainability of DL models has been the frontier of DL research at the present. It is crucial to make DL models more interpretative. Selvaraju et al., 2017 **[16]**, introduced the Grad-CAM technique or Gradient-weighted Class Activation Mapping, which is a powerful technique used in the field of DL offering a visual explanation of deep learning models.

DMRU has implemented Grad-CAM scoring scheme to pull out the genes from the transcriptome profile which appear to be critical for condition specific methylation for any given sequence stretch. This approach involves computing the gradient of a target class, denoted as "c," alongside the global average of the activation feature map "F" derived from the first convolutional layer "l." The gradient provides insight into how the model's performances change with respect to small variations in the input, allowing us to pinpoint features that are critical for distinguishing between different methylation states. By synthesizing these gradients with the activation maps, DMRU generates class-discriminative profiles which highlight the importance of specific features, expressing genes from the transcriptome profile in the present condition. To refine these profiles further, DMRU applies a weighted combination of the activation maps, effectively emphasizing the contribution of each feature based on its gradient score. Subsequently, a ReLU function is applied which ensures that only features with a positive influence are retained. This methodological framework allows us to visually represent and interpret which genes are responsible for the DNA sequence methylation states. The mathematical formulation and operational principles of Grad-CAM are described as follows:

$$GradCAM_l^c = ReLU\left(\sum_l \alpha_l^c F^l\right) \tag{11}$$

$$\alpha_l^c = \frac{1}{L}\sum_i \sum_j \left(\frac{\partial y^c}{\partial A_{ij}^l}\right) \tag{12}$$

Where, $\alpha_l^c$ denotes the neuron importance weights, highlighting the most relevant features that contribute to the models accuracy. ***'L'*** is the vector length in the feature map, and $A_{ij}^l$ represents the activation at **"*ij*$^{th}$"** position in the input array.

In the present implementation of Grad-CAM within DMRU, the first convolutional layer of the DenseNet architecture was selected as the layer of interest, as illustrated in **Figure 3**. This particular layer is pivotal because it captures the initial, low-level features of gene expression data, forming the foundation for more complex representations in subsequent layers. Following the methodology established by Selvaraju et al. (2017) **[16]**, Grad-CAM weighting technique was applied to quantify the significance of these gene expression representations. By performing a weighted summation of the distribution maps, an aggregated heatmap was derived that highlights the key genes implicated in DNA methylation activities. The results using the built models recapitulate multiple known trends. For example, the top-scoring genes corresponding to specific sequences frequently align with the documented gene ontology classifications related to the promoter sequences analyzed. This highlights the effectiveness of using Grad-CAM as a tool for elucidating the complexities of gene expression in the context of DNA methylation.

### Results and Discussion

#### Data collection and formation of datasets

DNA methylation has been reported to be a function of sequence properties and several genes **[17-29]**. Some of well identified such genes in plants are: DNA methyltransferase MET1, DDM1, VIM1–3, chromomethylase CMT1-3, histone methyltransferases kryptonite (KYP), Su(var)3-9 homologue 5 (SUVH5), IDN2, IDN2, DRM2, and SUVH6 **[17-22]**. However, these are usually fundamental genes which are not condition specific but usually expressed uniform way. While the fact is that in plants the regulatory DNA methylations are highly spatio-temporal, suggesting involvement of several other molecular factors in making the epigenetic regulation highly specific and time controlled.

In this study, we have investigated the relationship between DNA methylation profiles and transcriptome expression profiles through two important plant species, *Arabidopsis thaliana* and *Oryza sativa*. Our investigation was centered on the 2-kilobase (kb) upstream regions as the potential promoter regions where DNA methylations at cytosines have been reported critical in regulating the downstream gene's expression. The methylation profiles for all genes in both *A. thaliana* and *O. sativa* were recorded from various published studies. The methodology is thoroughly detailed in the methods section, with **Figure 2c** providing a visual representation of how these datasets were curated and utilized in this study. From the SRA database, a total of 260 RNA-seq fastq files and 232 WGBS-seq fastq files were populated for *A. thaliana* and *O. sativa*. This data encompassed a wide array of experimental conditions, 45 for *A. thaliana* and 40 for *O. sativa*. To ensure the integrity of the analysis, only those WGBS-seq data were included which had corresponding RNA-seq data for the same experimental conditions, allowing us to directly associate the gene expression levels with methylation status as well as build scope to consider the influence the of other genes too. For each gene, the 2kb upstream potential promoter sequences were extracted. These sequences were annotated for methylated cytosines, clearly demarcating methylated cytosines from the unmethylated ones. Each promoter sequence was represented multiple times, because of varying methylation states of cytosines across different conditions. In terms of scale, the dataset for *O. sativa* comprised 55,42,110 promoter sequences with variable methylation states, covering 38,756 genes and 40 conditions while *A. thaliana* contributed 51,14,144 promoter sequences with variable methylation states, covering 32,368 genes and 45 conditions. These data also used the corresponding gene expression data for the given condition from condition specific transcriptome profile. The average coverage of the RNA-seq data was ~17 for *A. thaliana* and ~8 for rice. These datasets have been named as Dataset “A” (*A. thaliana*) and Dataset “B” (*O. sativa*) in the current study, also are illustrated in **Figure 2** and details of the data provided in **Supplementary Table 2 Sheet 1-4**. A total of 45 experimental condition for *Arabidopsis* and 40 experimental condition for rice were taken.

#### Conjoint deep-learning on sequence properties and conditional transcriptome profiles raises an efficient model to detect cytosine methylation

The implementation of the DMRU model represents a significant advancement, particularly in the study of gene regulation through the interplay of DNA methylation, methylation region’s sequence information, and surrounding transcriptome’s profile including the expression of the downstream gene. As already mentioned above, the fundamental hypothesis was consideration of the fact that in plants DNA methylation is differential, where many other genes could be involved in the selection of condition specific cytosine methylation. Thus, a relationship between such expression pattern of causal genes, methylation spot’s local sequence information, and downstream gene’ expression may become a guide to evolve an efficient model where using the RNA-seq profile one may decode the DNA methylation pattern for the selected gene’s promoter region.

Central to this implementation is a composite deep co-learning system that leverages both Transformers and DenseNet architectures, which are adept at capturing complex patterns. This model was designed to understand the interdependent variability between methylation states in gene promoters and their corresponding RNA expression levels under varying environmental conditions. To rigorously evaluate the model's efficacy, the training and testing phases utilized Dataset “A”, having 70:30 split ratio for training and testing. It was ensured in this entire study that there was no overlap and redundancies existed between the training and testing sets, eliminating possibilities of any skews. The initial training phase involved 1,200 distinct promoter sequences paired with RNA-seq data across different experimental conditions. Complete implementation details of the deep-learning model is already covered in the methods section.

Once the dataset was processed through the integrated Transformers-DenseNet system, the model was initially evaluated on an unseen test set consisting of 200 promoter sequences across 45 experimental conditions covering a total of 31,600 different methylation profile sequences, which produced an impressive accuracy of 90.31% **(Figure 4a)**. This level of accuracy indicates that the model was successful in identifying methylated cytosines within the sequences, showcasing its potential for application in genomic studies to detect DNA methylation just using RNA-seq data. This system was further improved to obtain better results by conducting hyper-parameter optimization steps which is reported in the next section.

A very interesting now question arises that which features were most contributory in the rise of such accurate model to decode cytosine methylation. In this direction, ablation analysis was performed which evaluated one after another considered properties, and performance in their presence and in their absence. In the present section, we discuss the sequence based features' ablation analysis. A total of six distinct sequence-derived word representations were utilized: dimers, trimers, tetramers, pentamers, hexamers, and heptamers. Each representation corresponded to the number of consecutive nucleotides or characters considered within the sequences, thus providing varying lengths of contextual information. The underlying hypothesis was that integrating these representations would lead to additive effects that enhance the model's translational performance. The results indicated a clear trend: while individual word representations demonstrated limited capability most notably, the integration of these representations markedly improved performance due to enhanced sharing of contextual information **(Figure 4b)**. As more complex representations were introduced, significant increase in accuracy was observed: dimeric representations returned

26.02% accuracy, trimeric representation achieved 39.8%, tetrameric reached 57.13%, and pentameric representation attained 72.77%. This progression continued with hexameric and heptameric representations, which returned accuracy values of of 74% and 78%, respectively. It is evident from these findings that the contribution of higher-order words improved the performance mainly because of increase in the vocabulary due to higher number of representational words. But that came at the cost of computational complexity and memory requirements. For example, the GPU memory required for dinucleotides for the training set was 12 GB, which took 238 seconds of execuation time for single epoch. This shot up by manifolds when heptamers were considered where the required memory was 46 GB and took 1,456 seconds to complete the single epoch training process for the same set of training instances. Yet, it was very clear that the true potential of these representations was realized when they were combined. For instance, the pairing of pentameric and hexameric representations (totaling 3,991 words) resulted in a significant leap in the accuracy to 85.85%. Following this, the combination of hexamers with heptamers (3,989 words) further enhanced performance, yielding an accuracy of 89.32%. The best instance of the additive benefits of combining representations was illustrated when incorporating pentameric, hexameric, and heptameric representations, resulting in an impressive accuracy of 91.48% with a total of 5,985 representative words. These results emphasize the necessity of information sharing among different representations, as they collectively harnessed their unique attributes to raise a superior model. The findings are illustrated in performance plots **(Figure 4a)**, which clearly show the improvements in accuracy associated with various combinations of sequence representations.

#### Hyper-paramter optimization further improved the model

Above the model was developed with prefixed hyperparameters, though not completely random but values understood from our previous works on other problems **[8-10, 30]**. Initially, the main task was to determine if the identified sequence and expression properties were forming the right features or not. Additionally, the most suitable deep-learning architecture too had to be finalized. Once these were done and a substantial accuracy was obtained, hyperparamter optimization was the next essential to improve the performance of the developed deep-learning system.

An exploration into the performance of varying numbers of encoders and decoders for the Transformers-DenseNet system revealed a notable trend in the system's efficiency as layers were incrementally added. Specifically, the most pronounced increase in performance was observed at the addition of the eighth encoder-decoder pair, marking a significant point in the model's ability to handle the complexities of the task at hand. This enhancement in performance remained stable through the inclusion of subsequent layers up to the eighth, after which there was a noticeable decline in the effectiveness **(Figure 4c)**. In addition, an exploration into the performance of varying numbers of attention heads was also carried out with eight number of heads emerged as best for the model. The output from the multi-head attention layer was subjected to a dropout layer with a dropout fraction of 0.12, effectively regularizing the model to mitigate overfitting by randomly deactivating a fraction of the neurons during training. This output was then normalized to ensure that subsequent layers received data that was statistically stable and appropriately scaled. The architecture continued with a Feed-Forward network comprising 48 nodes with Relu activation function, followed by another dropout layer with a slightly higher dropout fraction of 0.14. This strategic layering approach allowed the model to not only learn complex representations but also to maintain a robust training regimen that promotes generalizability. Next, a second feed-forward layer was integrated, this time featuring 64 nodes with Selu activation function, further enriching the model's capacity for feature extraction and transformation. This layer was followed by another normalization stage, ensuring that the subsequent computations maintained a uniform scale and distribution. The output from the DenseNet component was then channeled through a feed-forward layer within the transformers' encoder, which played a crucial role in capturing the variability present in RNA expression profiles under specific conditions, along with their corresponding methylated sequence data. This architecture was designed to facilitate this interaction enabled the model to leverage rich biological data effectively, enhancing its performance capabilities. In terms of model training, the output from the encoder was transferred to the decoder's components through a masked multi-head attention mechanism, a design choice that crucially prevented information leakage during the training phase and ensured that the model adhered to the sequence generation rules intrinsic to natural language processing. This attention to detail culminated into the final processing via the decoder's feed-forward layer, responsible for generating coherent outputs in response to the processed inputs. To evaluate performance quantitatively, the model utilized the cross-entropy loss function for loss calculation, employing the Adam optimizer to enhance training efficiency and convergence speed. The learning rate for this optimizer was finally set at 0.001. The model underwent training across 100 epochs, with a batch size of 4 to optimize computational resources while maintaining the integrity of gradient updates. To prevent overfitting, we applied weight decay with a coefficient of 1e-5. This helped improve the performance of the model. We also implemented early stopping with a patience of 10 epochs, monitoring validation loss using PyTorch function ReduceLRonPlateau. Training was halted if the validation loss did not improve for 10 consecutive epochs, a standard practice, which helped prevent overfitting and reduce the training time. The hyperparameters defining the model's output layer were fine-tuned, resulting in a final configuration with softmax activation function, the cross-entropy loss function to guide learning, and the Adam optimizer. The “pt” format provided the model graph definition and weights to the pytorch structure while saving the model for translation. **Figure 3** illustrates the finally implemented deep-learning architecture. Upon evaluation against the test set, the optimized model achieved an impressive accuracy of approximately 93.17%, as illustrated in **Figure 4a**, a significant improvement by ~3% in accuracy exhibited by the optimized model after the hyperparameter tuning **(Figure 4a)**.

#### Remarkably consistent performance observed across wide range of experimental data

The investigation into the performance of the co-learning system when applied to the unseen promoter sequences yielded significant insights into its capabilities and performance. By evaluating upon 10,000 promoter sequences from Dataset "A" that were not included in the training and testing phases, we aimed to ascertain the model's generalizability beyond its initial training data. Remarkably, the model achieved an impressive accuracy of 92.82% **(as illustrated in Figure 4d)**, underscoring its effectiveness in capturing the intricate co-variability between RNA expression levels and the specific methylation preferences associated with these sequences in condition specific manner. This high level of accuracy suggests that the model learned well from the training data and internalized the underlying biological relationships, allowing it to make accurate discovery even when confronted with previously unseen data.

Following this promising result, we developed two co-learning model, Model "A" from Dataset "A" and Model "B" from Dataset "B". We further tested these models on the testset of Dataset "A" and Dataset "B", each comprising a total of 30,000 unique promoter sequences corresponding to *A. thaliana* and *O. sativa*, respectively. Both the model's performance remained consistent, with an average accuracy of 92.74% across the datasets. More specifically, the accuracy for *A. thaliana* was recorded at 92.6% from Model "A", while for *O. sativa* from Model "B", it slightly improved to 92.88% **(Figure 4d)**. This consistency in accuracy across datasets reinforces the model can reliably discover methylation patterns in diverse genomic and experimental conditions. **Figure 4f** provides the details of data splitting, subsequent stages in model development and noted performance leap. For detailed gene and condition wise metrics focusing on per-cytosine accuracy (includes accuracy, precision, recall, and F1-score alongside confusion matrices) specifically quantify how well the model translates unmethylated input sequences into correctly methylated output sequences, **Supplementary Table 3 Sheet 1** is provided. Afterwards, we assessed contribution from the integration of the RNA expression profile handled through DenseNet, the DL system was raised again without it. This analysis marked decrease in the model's accuracy, plummeting to 67.23% (p-value >> 0.01). Such a significant decline advocates for the necessity of jointly analyzing sequence data and RNA expression profiles which ultimately enriches the model's performance. 10-fold random trials of training and testing performance, raising models form randomly created training and testing data every time, concurred with the above observed performance level and scored in the same range consistently for Dataset "A". The MAE for training was found to be 0.0685, while for testing it was 0.0714, resulting in a very small difference of only 0.0029. A t-test comparing the MAE values for the training and test sets yielded a highly insignificant result of ~15%, much above the significance threshold of 5% or lower, further confirming the absence of any possibility of any significant over-fitting (**Supplementary Table S2 Sheet 5)**. For all these 10 cross-validations, the model consistently performed above 92% accuracy, suggesting a highly stable and consistent performance. **Figure 4g** provides the result of 10 fold random trails of train:test for both species. To conduct an unbiased performance testing without any potential recollection of data instances, it was ensured that no overlap and redundancy existed across any of the used data.

#### Accurate cross-species identification of DNA methylation where GC similarity is the key, not homology

Detecting DNA methylation computationally itself is a big challenge, whose next level comes with condition specific detection of DNA methylation. It it becomes top challenge when one looks for developing a tool which could work cross-species in total universal manner also. Universality is the requirement as any such tool should be capable to address any plant genome. Unfortunately, most of these challenges have remained unaddressed, which DMRU has successfully solved now.

The species specific models discussed above can tell condition specific methylation for any condition for the given species. But could they work with same efficiency on other species? In the context of cross-species application and universality of DMRU algorithm, the raised model was subjected to rigorous assessment to evaluate its effectiveness in identifying cross-species DNA methylation patterns. For this evaluation, we utilized two distinct datasets, referred to as Dataset "A" and Dataset "B," comprising sequences from two model plant, *A. thaliana* and *O. sativa*. The approach involved training a model using sequences from *A. thaliana* and subsequently testing it on sequences from *O. sativa*, while conversely, a model trained on *O. sativa* was tested on *A. thaliana* sequences. The model raised from *O. sativa* achieved a maximum accuracy of 83.9% (average accuracy 76.98%) on Dataset “A” covering *A. thaliana* **(Figure 4e)**. Similarly, the model raised from *A. thaliana* achieved a maximum accuracy of 79.89% (average accuracy 73.63%) on Dataset “B” **(Figure 4e)**. Both results clearly highlights that the raised models left enough scope for further improvement as they were far below than the accuracy levels observed for within species runs.

To develop a universal model capable of identifying DNA methylation promoter sequences from gene expression data across various species, our initial approach centered on leveraging homology between the sequences where the query sequences/genes were looked for their corresponding homologous ones from *A. thaliana.* This approach aimed to harness evolutionary relationships as has been the practice to expect similar functionality for homologous genes. However, despite rigorous testing across species, the model underperformed, achieving an accuracy of only around 83%, as depicted in **Figure 5a-b**. This outcome prompted us to rethink our approach as well as highlighted how wrong one can go while traditionally relying on the homology, especially for epigenetic controls. In epigenetic control it is GC isochore which should be looked for instead of homology. Therefore, the GC content (GC%) of the 2 Kb upstream regions for the genes were evaluated in both *A. thaliana and O. sativa*. By grouping genes with similar GC% value ranges, we created a new model based on these clusters. This innovative approach proved fruitful, when we applied the GC%-based model to the datasets from both species, the results were significantly improved. The model derived from *A. thaliana* demonstrated an impressive accuracy exceeding 90% on *O. sativa* (Dataset "B"), while the model based on *O. sativa* also achieved above 90% accuracy on *A. thaliana* (Dataset "A") **(Figure 5a-b)**. These outcomes underscore the critical role which GC content plays in the context of gene regulation and methylation identification.

In the next step, the above analysis was extended forward for the model's final assessment to determine how well it performed to identify cross-species DNA methylation on other unseen species. Based on experimental data availability for validation, this was done for *Arabidopsis lyrata* and *Solanum tuberosum*, for which WGBS-seq and RNA-seq data were available. Here also, the DMRU model developed from Dataset "A" achieved an average accuracy >90% for both the species, affirming its efficacy in handling cross-species conditions. **Figure 5c** provides the comparison details for both species by DMRU for cross-species performance. All these series of validation studies reinforced that DMRU performed consistently with high accuracy and emerged as a promising first of its kind tool for cross-species methylation identification without any dependency on predefined models while implementing generative and explainable AI innovative way. It can be employed successfully to even determine the methylated cytosines in newly sequenced plant genomes besides capturing the condition specific methylation profile with high accuracy and consistency.

#### DMRU outstands in comparative benchmarking studies

To evaluate the performance of our software tool in this study, we compared DMRU with nine different tools: iDNA-MS **[31]**, iDNA-ABT **[32]**, iDNA-ABF **[33],** MaskDNA-PGD **[34]**, MuLan-Methyl **[35]**, DeepPGD **[36]**, DeepSignal **[37],** PlantDeepMeth **[38],** and MethNet **[39],**. The four out of the compared nine tools viz. iDNA-MS, DeepPGD, MuLan-Methyl, and MethNet have bugs to run **(Suplementary Table 4 Sheet 1)**. Initially two different datasets based on *Arabidopsis thaliana* and *Oryza sativa*, were used to evaluate the performance of the developed model of DMRU with respect to five different tools, representing most recent and different approaches to detect DNA methylation: iDNA-ABT (BERT+ transductive information maximization (TIM)), MaskDNA-PGD (CNN, Bi-LSTM and attention mechanism), iDNA-ABF (BERT based approach), DeepSignal (CNN + BiLSTM), and PlantDeepMeth (CNN + Bi-GRU joint). The performance measure on the test set of these dataset which consists 12 different conditions gave an idea how the compared algorithms in their existing form perform. All these five tools were tested across these datasets where DMRU outperformed all of them, for all the performance metrics considered. All the five tools benchmarked here are not capable to identify condition specific DNA methylation and give a uniform cytosine methylation profile for all conditions, making them hardly of any use in real world applications. As highlighted in **Figure 6a-c and Supplementary Figure S3**, all these tools output exactly the same result and fail to capture the dynamic condition specific methylation patterns in *O. sativa*. On the other hand **(Figure 6d)**, DMRU captured the conditions specific methlyations in these regions. **Supplementary Figure S3-4** also presents dynamic condition specific methylation patterns in *A. thaliana.* **Supplementary Figure S5-7** show that the compared tools did not breach even 80% accuracy mark when the accuracy was calculated for every sequence under different conditions. With this all, DMRU stands at present the only approach which could capably suggest differential methylation patterns across different conditions, emerging also as a good alternative for costly bisulfite sequencing experiments.

AGO4) **(Figure 7e-h)**. Interestingly, in their study the authors had experimentally validated 12 genes and implicated them in Cuscuta induced stress, causing DNA methylation. All of these 12 genes were detected by DMRU Grad-CAM as influential genes associated with the observed DNA methyaltion patterns. All this reinforced the great credibility and utility of DMRU and its Grad-CAM to detect condition specific DNA methylation and infer associated influencer genes. This all also opens a plethora of opportunities and experimental interactions. The overlap between Grad-CAM genes and stress-responsive modules validates DMRU's ability to identify biologically relevant regulators of condition-specific methylation. For example:

**1) RNA-Directed DNA Methylation (RdDM):** Grad-CAM highlighted genes like PR1 (pathogenesis-related) and NLR (nucleotide-binding leucine-rich repeat receptors), which perceive Cuscuta-derived signals (e.g., CuRe1 peptides) and trigger RdDM components (NRPD1, RDR2, AGO4). This silences host defense genes or transposons via CHG/CHH hypermethylation, facilitating parasite nutrient uptake **[41-42]**.

**2) Mobile RNA exchange:** Cuscuta transfers mobile RNAs (mRNAs, siRNAs) to tomato, directing RdDM to alter host methylation patterns. Grad-CAM-identified genes like AGO4 (argonaute 4) and DRM2 (domains rearranged methyltransferase 2) are central to this process, mediating siRNA-guided DNA methylation **[43-44]**.

**3) Defense pathway modulation:** Enrichment in JA/SA signaling pathways aligns with reports that parasitism suppresses host immunity via methylation-dependent silencing **[45]**. For instance, hypermethylation of LOX2 (lipoxygenase 2) in susceptible tomatoes inhibits JA biosynthesis, reducing resistance **[46]**.

#### Web-server implementation

The DMRU server hosts the software in a user friendly manner for the identification of methylated cytosines within plant DNA sequences. To identify methylated cytosine, users are required to upload their DNA sequences in FASTA format and its associated experimental condition transcriptome expression profile in “tsv” format, followed by clicking the submit button. Once the user submits the data, the back end processing begins with the input data which is then passed through Transformer-DenseNet engine in the back-end which identifies the methylated cytosine for the given promoter sequences. DMRU server can handle upto 40 sequences in one submission. Subsequently, a results page featuring relevant information in an interactive format is presented. This result page consists of two tabs, **(i)** in the first tab, identified methyated cytosines were presented where the user can visualize the methylated cytosines within the given input sequences and also users get the capability to download the results in a tabular format, facilitating further downstream analysis and integration with other tools. **(ii)** The results page also provides a separate tab for explainability scoring line plot in the form of importance score while highlighting the genes important for the methylation in the sequence. This line plot is implemented through plotly.js. Users can download results in a tabular format. The server implementation has been done using PHP, JavaScript, HTML5, and the Apache-Linux platform. The majority of codes were developed in Shell and Python scripts for jobs related to data curation and back-end processing. The project was executed within the Ubuntu Linux platform's Open Source OS environment. **Figure 8** provides an overview of DMRU webserver implementation. The DMRU system is freely accessible at https://scbb.ihbt.res.in/DMRU/.

### Pseudocode of the Transformer-DenseNet model architecture

**DenseNet-121**

**//input_tensor: 3D tensor [300 X 36 X 3] for DenseNet121 each value in this tensor represents a gene with expression values**

```
function DenseNet121(input_tensor):
{
//   Applies DenseNet-121 to the input 3D gene expression tensor.
//   Parameters:
//       input_tensor: Tensor of shape [batch_size x channels x height x width]
//   Returns:
//       features: Tensor of shape [sequence_length x d_densenet]
    features = CNN_ForwardPass(input_tensor)  # Apply pre-trained or custom DenseNet-121
    return features
}

function TransformerEncoderWithDenseNet(X_kmers, cnn_input, N_layers, num_heads, d_model, d_ff):
{
//    Combines k-mer token embeddings with CNN features via DenseNet-121, processes through Transformer encoder.
//   Parameters:
//       X_kmers: List of k-mer tokens (e.g., ["ATC", "TCG", ...])
//       cnn_input: 3D tensor for DenseNet feature extraction
//       N_layers: Number of Transformer encoder layers
//       num_heads: Number of self-attention heads
//       d_model: Dimensionality of model (embedding size)
//       d_ff: Hidden size of feed-forward layers
```

```
//    Returns:
//        encoder_output: Tensor of shape [seq_len x d_model]

//   Step 1: Token Embedding + Positional Encoding
    X = EmbedTokens(X_kmers)                              # → [seq_len x d_model]
    X += PositionalEncoding(length=len(X), d_model=d_model) # Add positional information

//    Step 2: Extract CNN features from DenseNet
    cnn_features = DenseNet121(cnn_input)                  # → [seq_len x d_densenet]

//    Step 3: Transformer Encoder Layers
    for layer in range(N_layers):
    do
    {
//  Multi-Head Self-Attention
        Q = X @ W_Q
        K = X @ W_K
        V = X @ W_V

        Q_heads = split_into_heads(Q, num_heads)           # → [num_heads x seq_len x d_k]
        K_heads = split_into_heads(K, num_heads)
        V_heads = split_into_heads(V, num_heads)
        attention_heads = []
        for h in range(num_heads):
         do
```

```
        {
          scores = (Q_heads[h] @ transpose(K_heads[h])) / sqrt(d_k)
          weights = softmax(scores)
          head_output = weights @ V_heads[h]
          attention_heads.append(head_output)
        }
      concatenated = concat_heads(attention_heads)        # → [seq_len x d_model]
      attention_output = concatenated @ W_O               # Linear projection of attention output

//    Residual + Normalization
      X = LayerNorm(X + attention_output)

//    Feed-Forward + CNN Feature Fusion
      FFN_output = ReLU(X @ W1 + b1)                      # → [seq_len x d_ff]

//     Concatenate with DenseNet features
      fusion = concat([FFN_output, cnn_features], axis=-1)  # → [seq_len x (d_ff + d_densenet)]

//     Optional projection to match d_model
      fused_output = fusion @ W_fusion + b_fusion         # → [seq_len x d_model]

//     Residual + Normalization
      X = LayerNorm(X + fused_output)
    }
  encoder_output = X
```

```
    return encoder_output  # shape: [seq_len x d_model]
}

function TransformerDecoder(Y_input_tokens, encoder_output, N_layers, num_heads, d_model,
d_ff):
{
 //   Standard Transformer decoder that uses encoder output for cross-attention.
//    Parameters:
//        Y_input_tokens: Target sequence tokens (decoder input)
//        encoder_output: Output from the encoder [seq_len x d_model]
//    Returns:
//        decoder_output: Final decoder hidden states [seq_len x d_model]

// Step 1: Token Embedding + Positional Encoding
    Y = EmbedTokens(Y_input_tokens)                    # → [seq_len x d_model]
    Y += PositionalEncoding(length=len(Y), d_model=d_model)

    for layer in range(N_layers):
    do
    {
//      Masked Multi-Head Self-Attention (causal)
        Q = Y @ W_Q_self
        K = Y @ W_K_self
        V = Y @ W_V_self
        Q_heads = split_into_heads(Q, num_heads)
```

```
    K_heads = split_into_heads(K, num_heads)
    V_heads = split_into_heads(V, num_heads)
    masked_attention_heads = []
    for h in range(num_heads):
    do
    {
        scores = (Q_heads[h] @ transpose(K_heads[h])) / sqrt(d_k)
        scores = apply_causal_mask(scores)          # Prevent future attention means applying a
// causal mask so each decoder step attends only to current and past tokens, not future ones.
        weights = softmax(scores)
        head_output = weights @ V_heads[h]
        masked_attention_heads.append(head_output)
        }
    self_attn_output = concat_heads(masked_attention_heads) @ W_O_self
    Y = LayerNorm(Y + self_attn_output)

//     Cross-Attention with Encoder Output
    Q = Y @ W_Q_cross
    K = encoder_output @ W_K_cross
    V = encoder_output @ W_V_cross
    Q_heads = split_into_heads(Q, num_heads)
    K_heads = split_into_heads(K, num_heads)
    V_heads = split_into_heads(V, num_heads)
    cross_attention_heads = []
    for h in range(num_heads):
```

```
        do
        {
            scores = (Q_heads[h] @ transpose(K_heads[h])) / sqrt(d_k)
            weights = softmax(scores)
            head_output = weights @ V_heads[h]
            cross_attention_heads.append(head_output)
         }
        cross_attn_output = concat_heads(cross_attention_heads) @ W_O_cross
        Y = LayerNorm(Y + cross_attn_output)

//      Feed-Forward Network
        FFN_output = ReLU(Y @ W1 + b1)
        FFN_output = FFN_output @ W2 + b2
        Y = LayerNorm(Y + FFN_output)
    }
    decoder_output = Y
    return decoder_output  # → [seq_len x d_model]
}

function GenerateOutput(decoder_output, vocab_size):
{
//    Maps decoder output to vocabulary probabilities.
//    Parameters:
//        decoder_output: Output of the decoder [seq_len x d_model]
//        vocab_size: Size of target vocabulary
```

```
//    Returns:
//        output_probs: Probability distribution over vocabulary [seq_len x vocab_size]
    logits = decoder_output @ W_vocab + b_vocab  # Linear projection
    output_probs = softmax(logits, axis=-1)
    return output_probs
}
```

#### Supplementary data

**Supplementary Table S1: Brief list of some of the published tools for DNA methylation identification.**

**Supplementary Table S2: Information regarding to (A)** dataset "A" and “B” general information, **(B)** Hyperparameter optimization and details of Transformers, **(C)** Details of abylation and tenfold MAE analysis, **(D)** sRNA-seq, **(D)** Statistics of the training curves (loss/accuracy/gpu usage vs. epochs) during training and testing process.

**Supplementary Table S3: Statistics of the key metrics such as precision, recall, or F1-score are shown across all test conditions.**

**Supplementary Table S4: Statistics of the bugs identified during benchmarking and data regarding GC% and methylation content in transcription factor binding sites.**

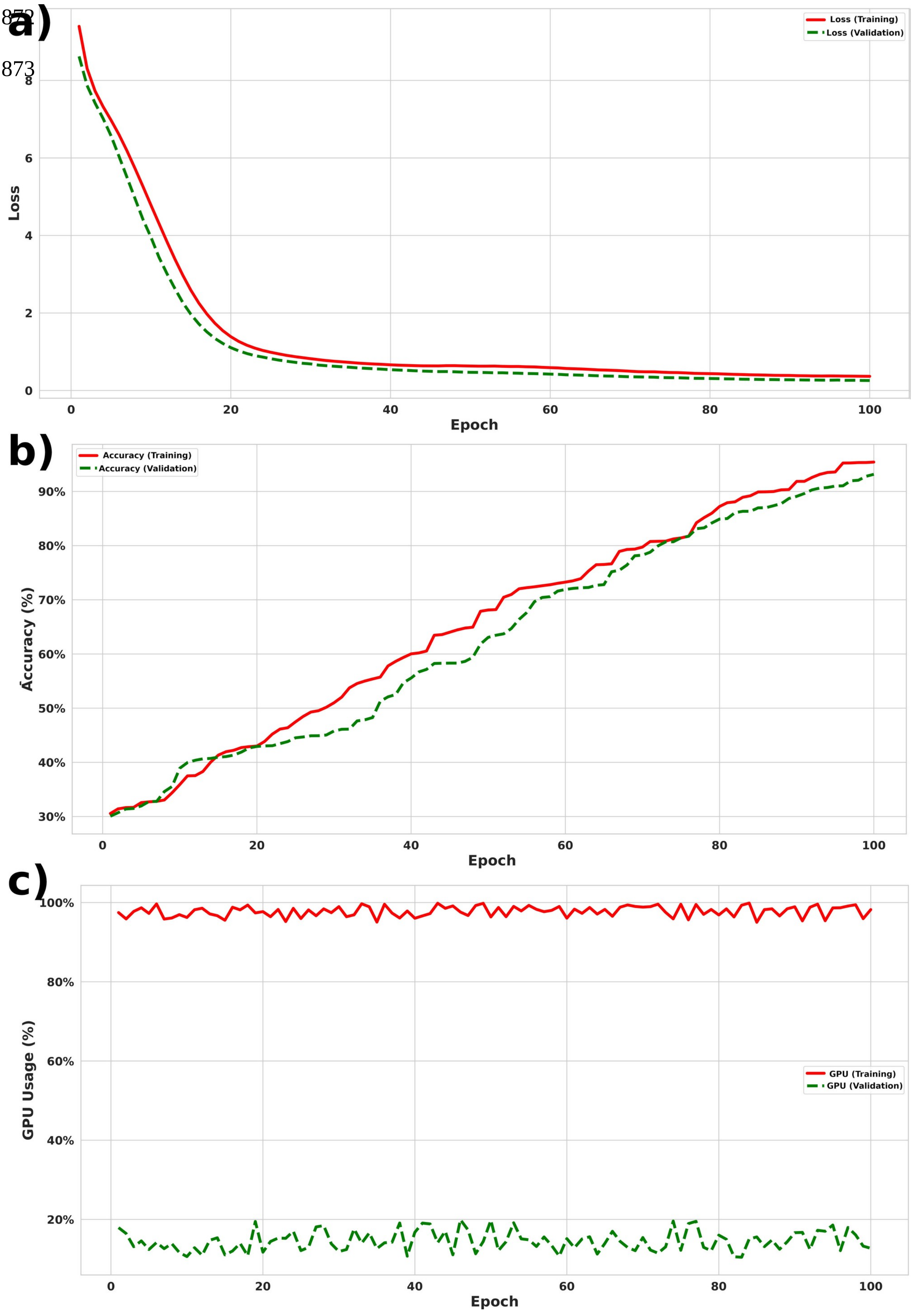
a)
Loss (Training)
Loss (Validation)
Loss
Epoch
b)
Accuracy (Training)
Accuracy (Validation)
Accuracy (%)
Epoch
c)
GPU (Training)
GPU (Validation)
GPU Usage (%)
Epoch

**Supplementary Figure S1: Training and validation loss, accuracy, and GPU usage over epochs.** **a)** This line plot illustrates the loss progression during training and validation phases across epochs. The training loss is shown in solid red, and the validation loss is shown in dashed green. The loss values are tracked to evaluate model convergence and potential overfitting. **b)** This figure shows the training and validation accuracy (%) across epochs. Accuracy values are displayed as percentages, reflecting the model's classification performance during each training epoch. **c)** This plot depicts GPU utilization (%) during training and validation over time.

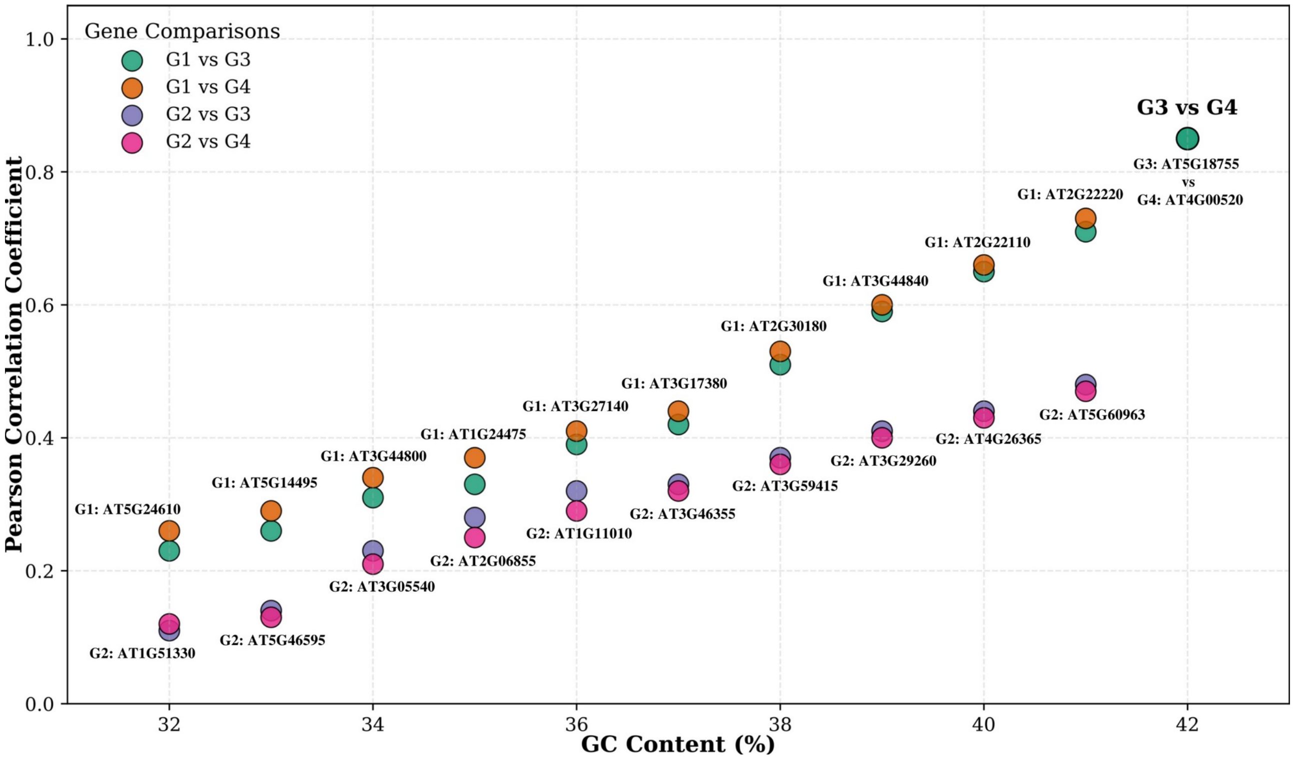


**Supplementary Figure S2: Two randomly selected genes from GC% group of 42% GC content were initially selected (G3:AT5G18755, G4:AT4G00520).** Within group, we observed high expression correlation between the member genes. These two genes too displayed the same (r ≈ 0.85). When these genes were compared with other GC% groups, the expression similarity gradually vanished as the GC dissimilarity increase. In the present figure we randomly selected two representative genes for each GC% group.

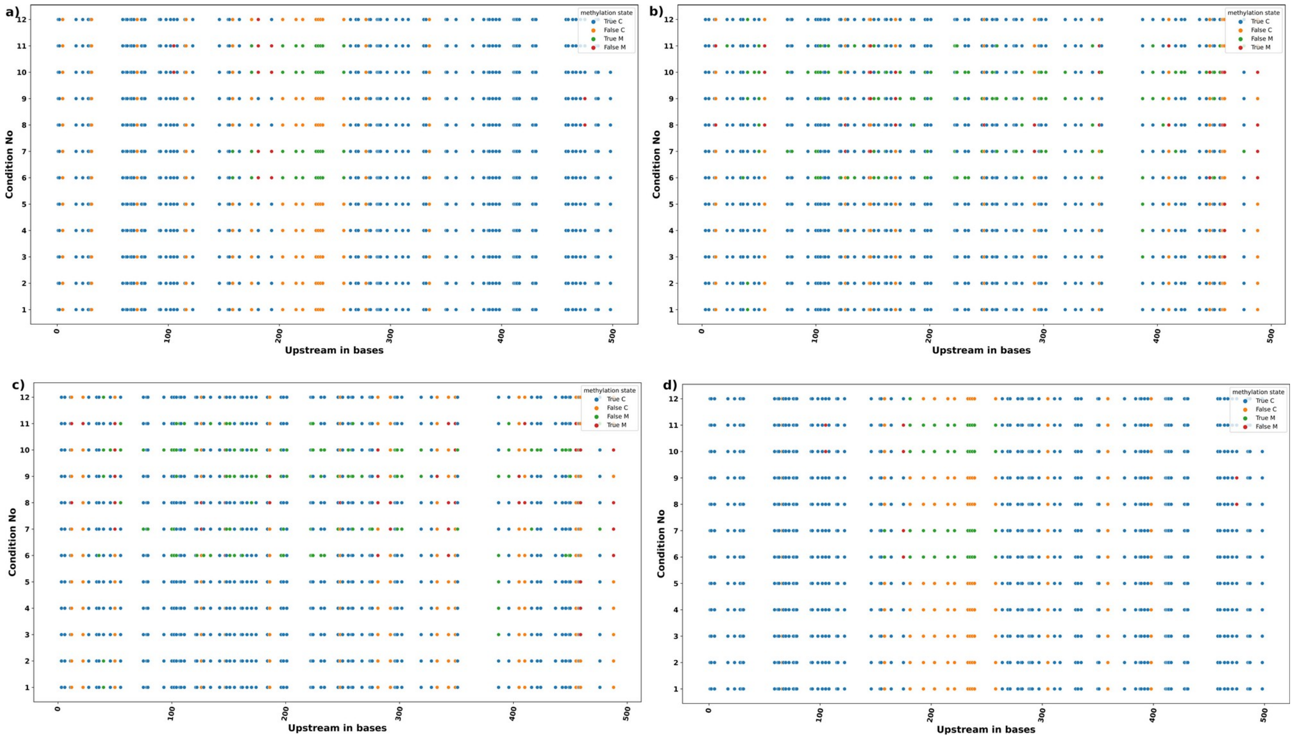


**Supplementary Figure S3: The performance over the 12 different experimental conditions for the compared software tools were benchmarked for the upstream region (have 500 bases shown) of gene AT1G01010 and Os01g0110100, for the tools (a, c) DeepSignal, and (b, d) PlantDeepMeth.**

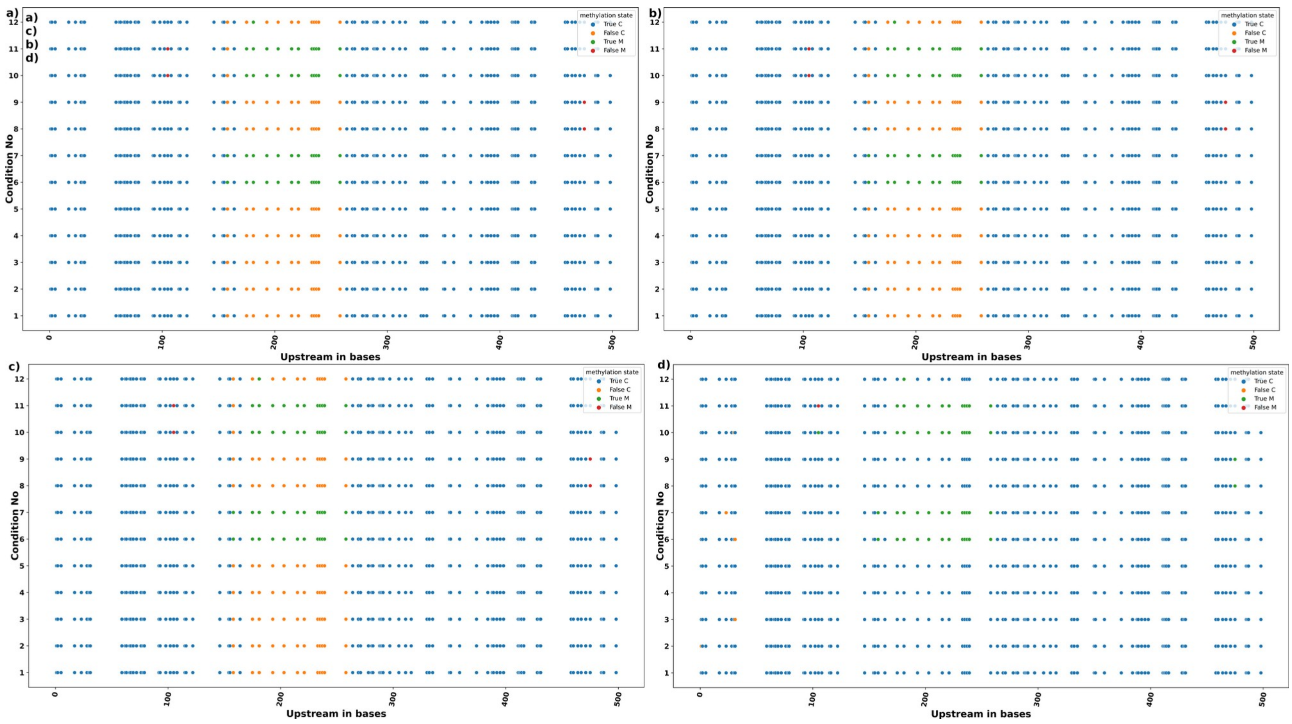


**Supplementary Figure S4: Performance and benchmarking of the universal model of DMRU based on GC% for *Arabidopsis thaliana*.** The performance over the 12 different experimental conditions for the compared software tools were benchmarked for the upstream region (have 500 bases shown) of gene AT1G01010, for the tools **(A)** iDNA-ABF, **(B)** iDNA-ABT, **(C)** MaskDNA-PGD, and **(D)** DMRU.

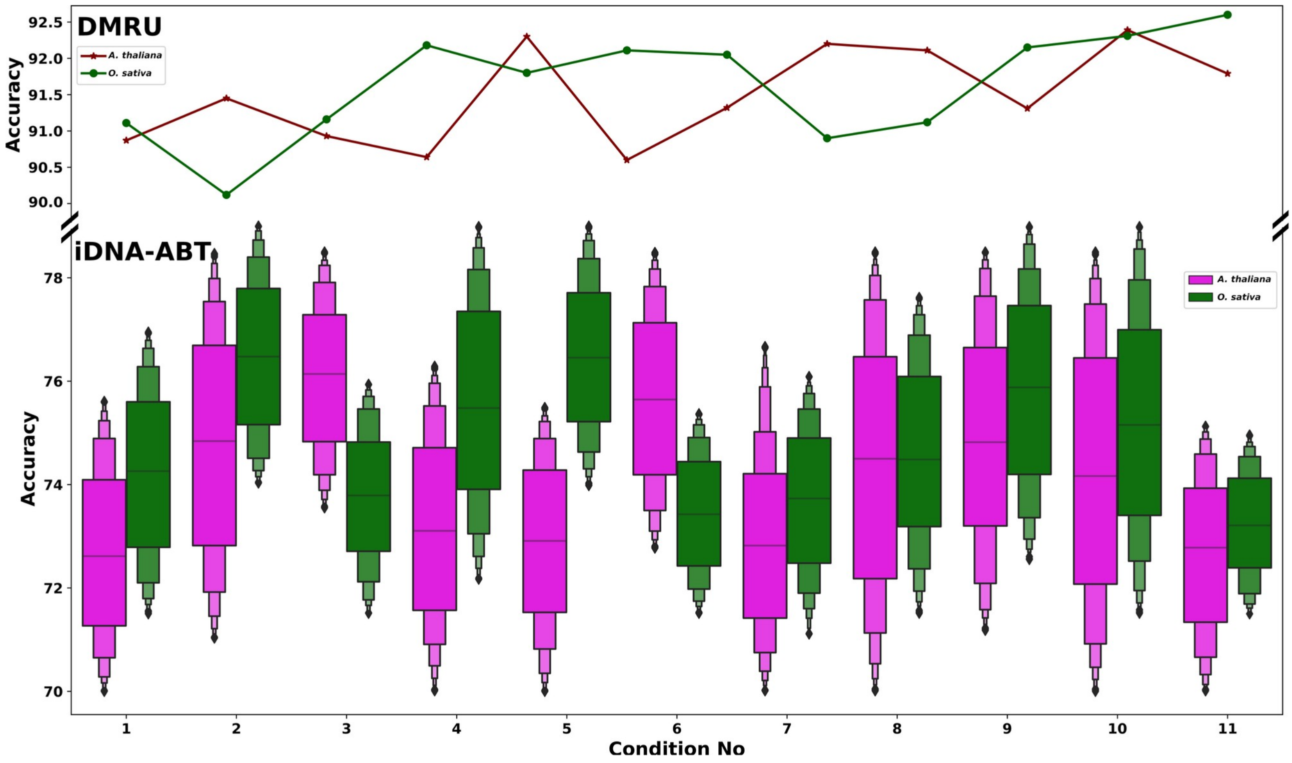


**Supplementary Figure S5: Performance accuracy of the benchmarked software tools (DMRU and iDNA-ABT) across different conditions.** iDNA-ABT did not breach even 80% accuracy mark while DMRU easily crosses 90% when the accuracy was calculated for every sequence under different conditions. Rendering DMRU effective across different experimental conditions.

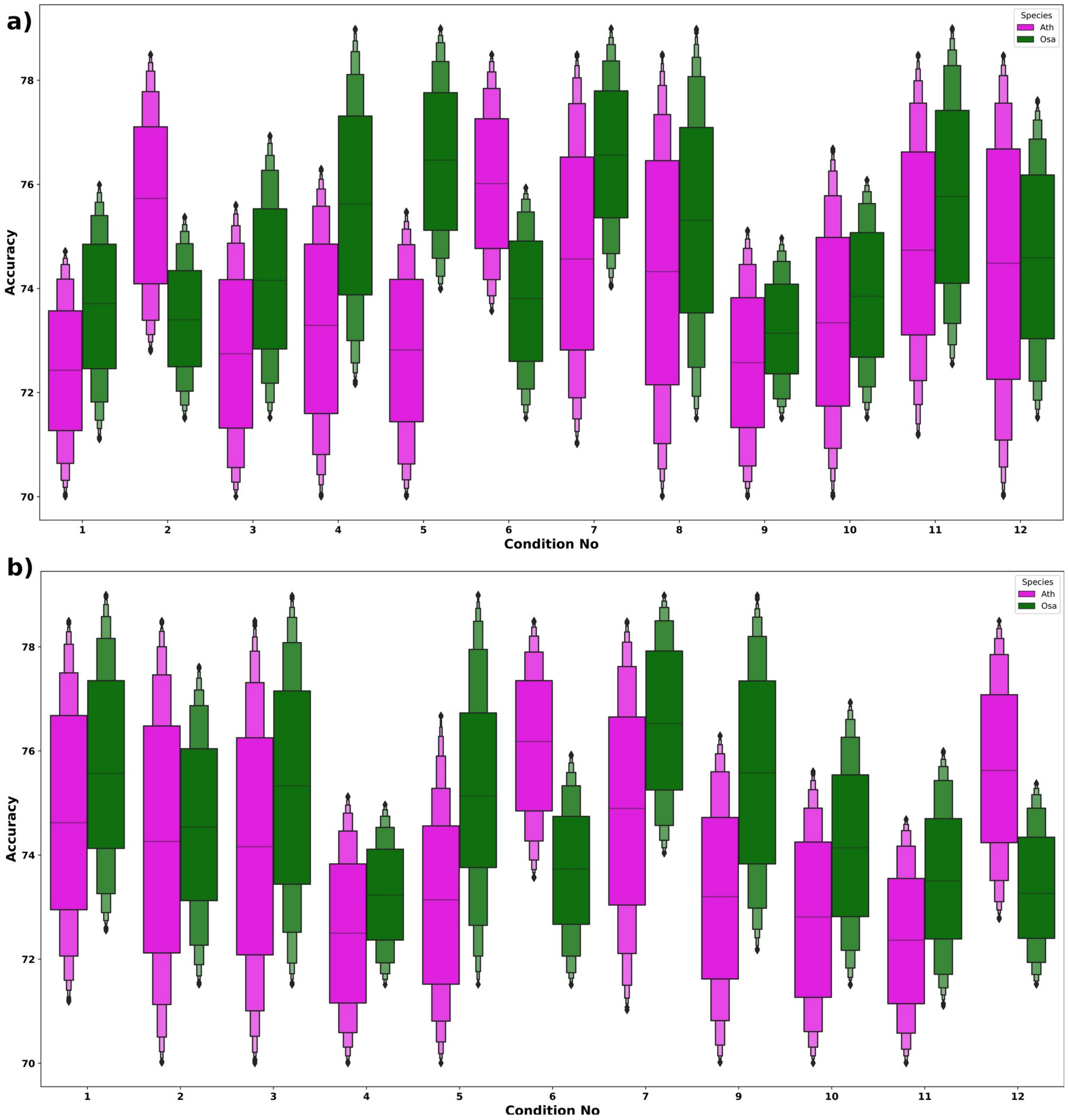


**Supplementary Figure S6: Performance accuracy of the software tools across different conditions.** **(A)** iDNA-ABF and **(B)** MaskDNA-PGD. These software tools did not breach even 80% accuracy mark when the accuracy was calculated for every sequence under different conditions.

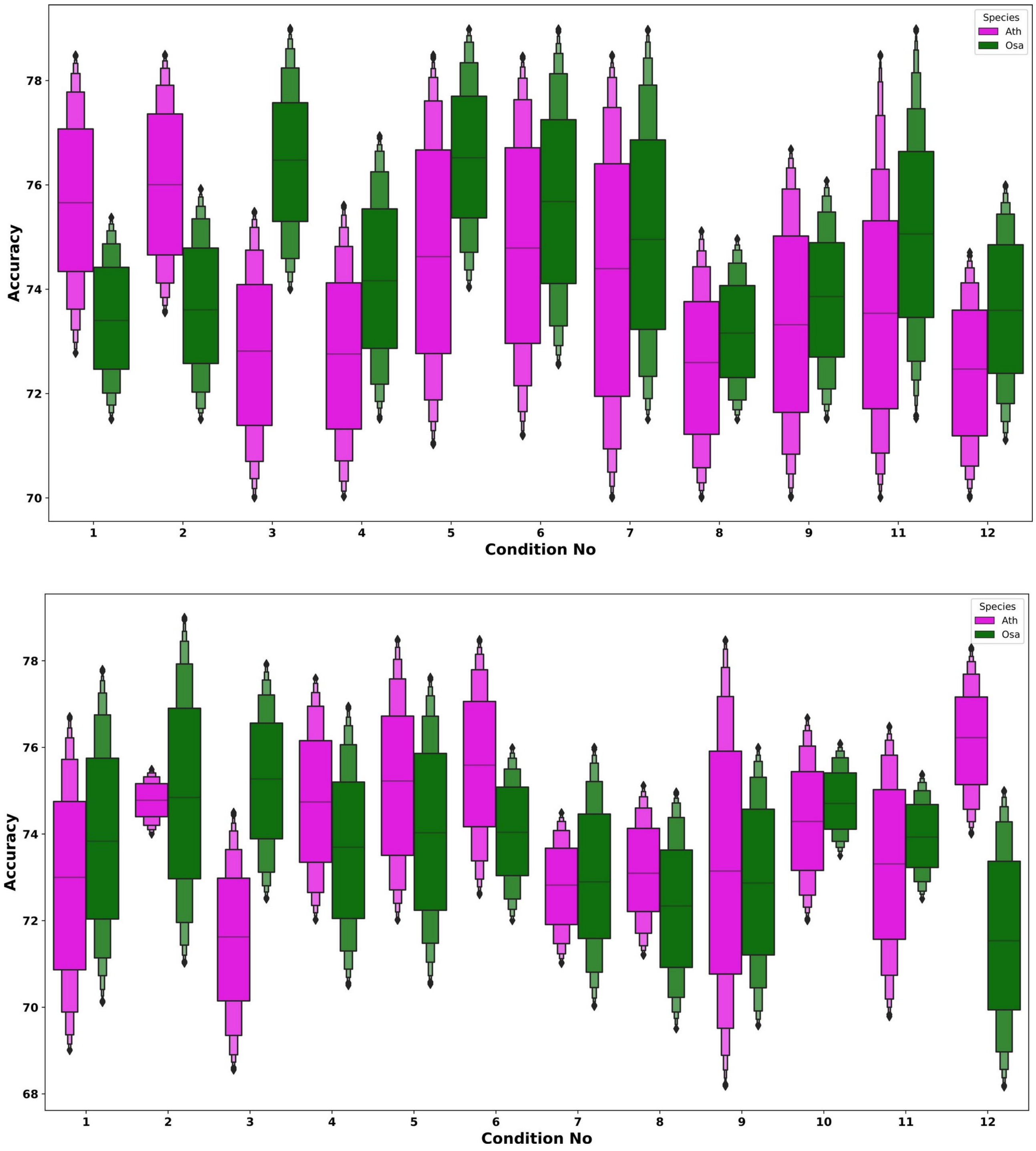


**Supplementary Figure S7: Performance accuracy of the software tools across different conditions. (A)** DeepSignal and **(B)** PlantDeepMeth. These software tools did not breach even 80% accuracy mark when the accuracy was calculated for every sequence under different conditions.

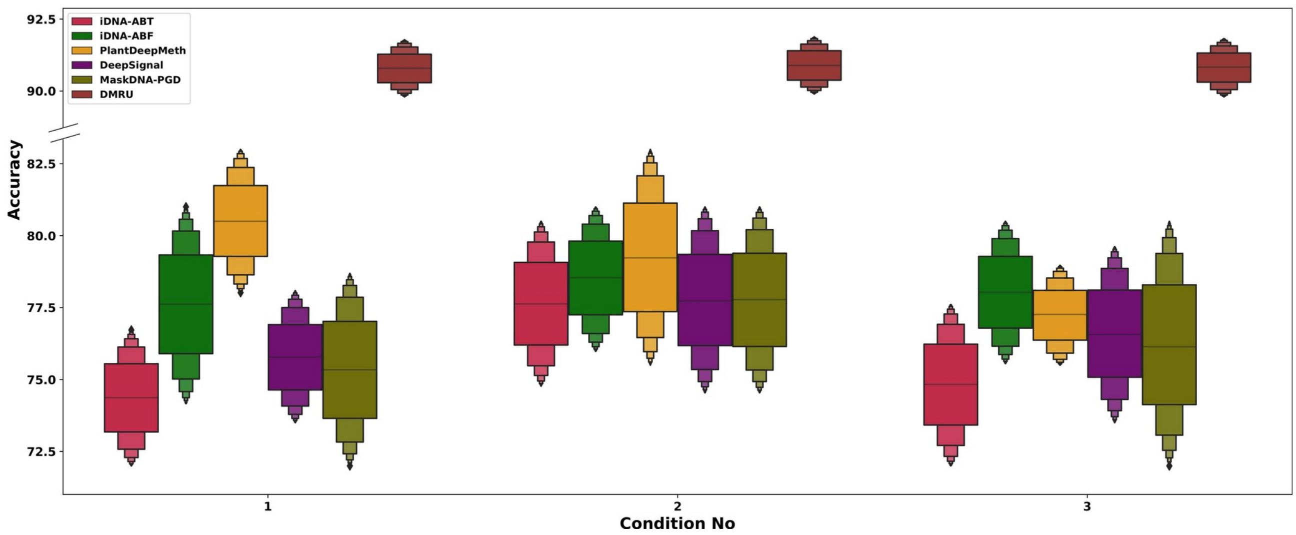


**Supplementary Figure S8: Performance accuracy of the software tools across different conditions on *Zea mays*.** Software tools other than DMRU did not breach even 80% accuracy mark when the accuracy was calculated for every sequence under different conditions.

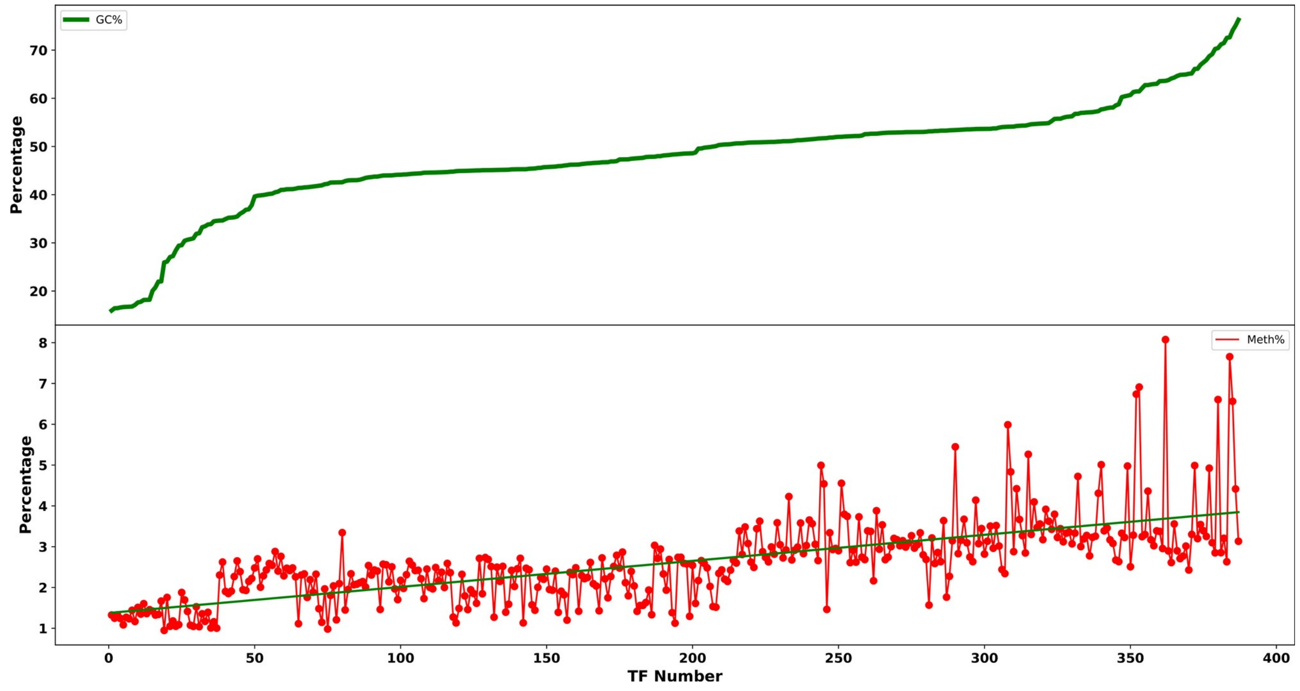


**Supplementary Figure S9: Comparison of GC content and methylation levels across transcription factors binding sites (TFBSs).** The line plot shows the GC percentage distribution (GC%) of TFBS sequences in *Arabidopsis thaliana*. The scatter plot displays the corresponding methylation percentage for each TF (in red). A green regression line indicates the overall pattern of methylation changes with increasing GC%. Higher GC is mostly associated with higher chance of DNA methylation.

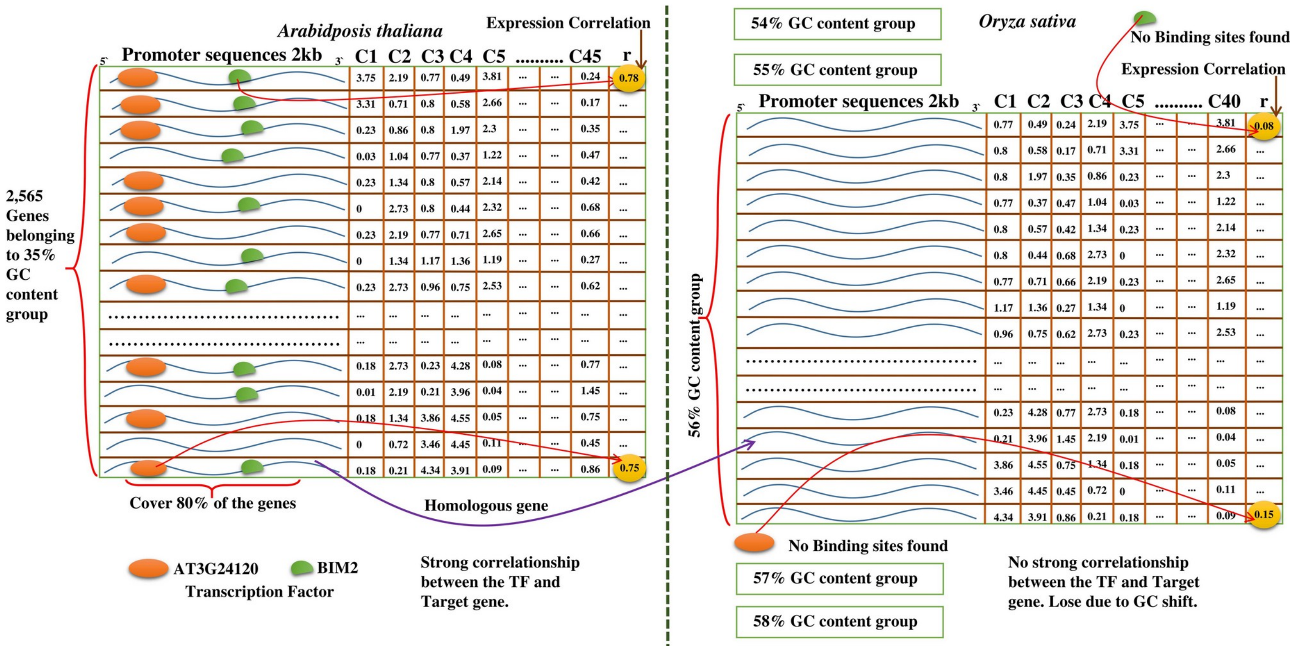


**Supplementary Figure S10: Comparative analysis of transcription factor (TF) binding and expression correlation in genes grouped by GC content in *Arabidopsis thaliana* and *Oryza sativa*.** The left side represents genes from *A. thaliana* belonging to the 35% GC content group. Promoter regions (2 kb upstream) of these genes were analyzed for TF binding sites, revealing that AT3G24120 and BIM2 together cover over 80% of the gene set. Expression correlation (Pearson's r) between TFs and selected genes within the group is shown in the matrix, with strong positive correlations observed, indicating strong co-regulation. The right side shows the analysis in *O. sativa*, where the homologous gene (OsNAC06) belongs to the 56% GC content group. Promoter analysis of this group did not identify any binding sites for the homologous TFs. Corresponding expression correlation between these TFs and selected target genes including (OsNAC06) shows no strong correlation, suggesting a loss of regulatory interaction likely due to the GC content shift across the compared species.
