## Supplemental Table S1 for "DMRU: Generative Deep-Learning to unravel condition specific cytosine methylation in plants"

**Tables**

**Supplementary Table S1: Brief list of some of the published tools for DNA methylation identification.**

| S.No. | Software | Algorithm | Encoding scheme | Methylation Type | Dataset | Species | Year | Webserver (W)/ Standalone (S) |
| --- | --- | --- | --- | --- | --- | --- | --- | --- |
| 1 | IDNA4mC [1] | Support Vector Machine (SVM) | Nucleotide chemical properties (NCP) and nucleotide frequency (NF) | N^4^-methylcytosine | Caenorhabditis elegans, Drosophila melanogaster, Arabidopsis thaliana, Escherichia coli, Geoalkalibacter subterraneus and Geobacter pickeringii | Animal & Plant | 2017 | W |
| 2 | 4mCPred [2] | SVM | Position-specific trinucleotide propensity (PSTNP) and electron-ion interaction potential (EIIP) | N^4^-methylcytosine | *C.* elegans, D. melanogaster, A. thaliana, E. coli, G. subterraneus and G. pickeringii | Animal & Plant | 2019 | S |
| 3 | i6mA-Pred [3] | SVM | NCP and NF | N^6^-methyladenine | Oryza sativa | Plants | 2019 | S |
| 4 | MM-6mAPred [4] | Markov algorithm | Transition probability between adjacent nucleotides | N^6^-methyladenine | Oryza sativa | Plants | 2020 | S |
| 5 | 4mcDeep-CBI [5] | Convolutional Neural Netowrk (CNN) & Biderectional Long short Term Memory (BiLSTM) | Binary and k-mer frequency (BKF), Dinucleotide binary profile and frequency (DBPF), K-Nearest Neighbor (KNN), Physical-Chemical Properties (PCP), Multivariate Mutual Information (MMI), Pseudo dinucleotide composition (PseDNC), Electron-ion interaction pseudopotentials of trinucleotide (PseEIIP), and Ring-function-hydrogen-chemical properties (RFHCP) | N^4^-methylcytosine | C. elegans | Animal | 2020 | S |
| 6 | iDNA-MS [6] | Random forest (RF) | *K*-tuple nucleotide component (KTNC), NCP, NF, and mono-nucleotide binary encoding (MNBE) | DNA methylation | C. equisetifoli, Fragaria vesca, S. cerevisiae, A. thaliana, C. elegans, D. melanogaster, Homo sapiens, Rosa chinensis, Tolypocladium, Tetrahymena thermophile, Xanthomonas oryzae PV. Oryzicola, and Mus musculus | Animal & Plant | 2020 | W/S |
| 7 | 6mA-Finder [7] | RF | Accumulated Nucleotide Frequency (ANF), Binary, Composition of K-spaced Nucleic Acid Pairs (CKSNAP), Dinucleotide Composition (DNC), Enhanced Nucleic Acid Composition (ENAC), Nucleic Acid Composition (NAC) and Trinucleotide Composition (TNC), EIIP, NCP, and PseDNC | N^6^-methyladenine | Mus musculus | Animal | 2020 | S |
| 8 | DNC4mC-Deep [8] | CNN | 2Kmer, 3Kmer, binary encoding (BE), NCP, NF, and MMI | N^4^-methylcytosine | F. vesca, R. chinensis | Plants | 2020 | W |
| 9 | iDNA-MT [9] | Bidirectional gated recurrent units (BiGRU) | Dimer word embeddings | N^4^-methylcytosine and N^6^-methyladenine | C. equisetifolia, F. vesca, S. cerevisiae, and Tolypocladium | Animal & Plant | 2021 | NA |
| 10 | Deep4mC [10] | Deep CNNs with attention mechanism | ANF, Binary, CKSNAP, PseEIIP, NAC, DNC, TNC, ENAC, Kmer, Reverse compliment Kmer (RCK), NCP, PseDNC | N^4^-methylcytosine | A. thaliana, C. elegans, D. melanogaster, E. coli, G. pickeringii, G. subterraneus | Animal & Plant | 2021 | W |
| 11 | DeepTorrent [11] | CNNs with inception, BiLSTM and an attention layer | One hot encoding (OHE) | N^4^-methylcytosine | C. elegans, D. melanogaster, A. thaliana, E. coli, G. subterraneus and G. pickeringii | Animal & Plant | 2021 | S/W |
| 12 | iDNA-ABT [12] | Bidirectional encoder representations from transformers (BERT) together with transductive information maximization (TIM) | Embedding | N^4^-methylcytosine, N^5^-methylcytosine, and N^6^-methyladenine | H. sapiens, M. musculus, A. thaliana,, C. elegans,, Casuarina equisetifolia,, D. melanogaster, F. vesca, R. chinensis, S. cerevisiae, Tolypocladium sp SUP5-1, Tetrahymena thermophile, and Xanthomonas oryzae | Animal & Plant | 2021 | S/W |
| 13 | Deep-4mCGP [13] | CNN | OHE | N^4^-methylcytosine | *G. pickeringii* | Bacteria | 2022 | S |
| 14 | i6mA-Vote [14] | Integrating RF, Multi Layer Perceptron (MLP), Stochastic gradient descent (SGD), linear discriminant analysis (LDA), extreme gradient boosting (XGB), based on majority voting strategy | Dinucleotide OHE | N^6^-methyladenine | *Rosa*, Rice, and *Arabidopsis* | Plants | 2022 | S |
| 15 | MGF6mARice [15] | CNN | Simplified molecular input line entry system (SMILES) | N^6^-methyladenine | Rice | Plant | 2022 | S |
| 16 | Mouse4mC-BGRU [16] | BiGRU | Embedding | N^4^-methylcytosine | Mus musculus | Animal | 2022 | NA |
| 17 | Deep6mAPred [17] | CNN & BiLSTM | OHE | N^6^-methyladenine | Rice, *F. Versa, R. chinensis* | Plant | 2022 | W |
| 18 | bert-DNA [18] | Transformer & CNN | Word Embedding | N^6^-methyladenine | Mus musculus | Animal | 2022 | S |
| 19 | iDNA-ABF [19] | BERT | 6mer, 3mer Word embedding | N^4^-methylcytosine, N^5^-methylcytosine, and N^6^-methyladenine | C. equisetifoli, F. vesca, S. cerevisiae, A. thaliana, C. elegans, D. melanogaster, H. sapiens, R. chinensis, Tolypocladium, T. thermophile, Xanthomonas oryzae PV. Oryzicola, and M. musculus | Animal & Plant | 2022 | S/W |
| 20 | MaskDNA-PGD [20] | CNN, BiLSTM and attention mechanism | Word embedding | N^4^-methylcytosine, N^5^-methylcytosine, and N^6^-methyladenine | C. equisetifolia, F. vesca, S. cerevisiae. H. sapiens, A. thaliana, C. elegans, D. melanogaster, Tolypocladium, and T. thermophile | Animal & Plant | 11, 2022 | S |
| 21 | MuLan-Methyl [21] | Transformers | Word embedding | N^4^-methylcytosine, N^5^-methylcytosine, and N^6^-methyladenine | *T. thermophile, A. thaliana, C. elegans, D. melanogaster, H. sapiens, C. equisetifoli, F. vesca, S. cerevisiae, R. chinensis, Tolypocladium, T. thermophile, and M. musculus* | Animal & Plant | 12, 2023 | S/W |
| 22 | i5mC-DCGA  **[22]** | Convolutional Block Attention Module (CBAM), BiGRU and attention mechanism | OHE | N^5^-methylcytosine | Human | Animal | 2024 | S |
| 23 | DeepPGD  [23] | Temporal Convolution, BiLSTM, and Attention Mechanism | 3mer word embedding | N^4^-methylcytosine, N^5^-methylcytosine, and N^6^-methyladenine | C. equisetifolia, F. vesca, S. cerevisiae. H. sapiens, A. thaliana, C. elegans, D. melanogaster, Tolypocladium, and T. thermophile | Animal & Plant | 2024 | S |
| 24 | MethBERT  [24] | Transformer (BERT) | Learnable positional embeddings + ionic signals from Oxford Nanopore | 5mC (CpG) | E. coli and human NA12878 (Nanopore R9) | Bacteria, Human | 2021 | S |
| 25 | CpG-transformer [25] | Transformer (axial + sliding-window self-attention) | CpG methylation matrices from single-cell BS-seq/scRRBS-seq | CpG (single-cell) | Human single-cell datasets | Human single-cells | 2022 | S |
| 26 | DeepSignal [26] | CNN + BiLSTM | Raw ionic Nanopore signals + sequence features | 5mC and 6mA (CpG & adenine) | Human (NA12878), E. coli, pUC19 | Bacterial & Human | 2019 | S |
| 27 | PlantDeepMeth [27] | CNN + Bi-GRU joint model | DNA sequence + neighboring methylation context | CpG, CHG, CHH (plant 5mC) | Brassica rapa and Arabidopsis thaliana | Plant | 2021 | S |
| 28 | MethNet [28] | Elastic-net regression + cis-regulatory network modeling | CpG site methylation (TCGA array/WGBS); gene expression; cis-distance weighting; integration across multi-cancer samples | CpG methylation (5mC) | Large-scale TCGA methylation & expression across multiple cancers; validated with promoter capture Hi-C & CRISPRi Perturb-seq | Human (multiple cancers) | 2024 | S |
